# Allosteric mechanism of signal transduction in the two-component system histidine kinase PhoQ

**DOI:** 10.1101/2021.09.03.458835

**Authors:** Bruk Mensa, Nicholas F. Polizzi, Kathleen Molnar, Andrew Natale, Thomas Lemmin, William F. DeGrado

## Abstract

Transmembrane signaling proteins couple extracytosolic sensors to cytosolic effectors. Here, we examine how binding of Mg^2+^ to the sensor domain of an *E. coli* two component histidine kinase (HK), PhoQ, modulates its cytoplasmic kinase domain. We use cysteine-crosslinking and reporter-gene assays to simultaneously and independently probe the signaling state of PhoQ’s sensor and autokinase domains in a set of over 30 mutants. Strikingly, conservative single-site mutants distant from the sensor or catalytic site strongly influence PhoQ’s ligand-sensitivity as well as the magnitude and direction of the signal, endowing diverse signaling characteristics without need for epistasis. Data from 35 mutants are explained by a semi-empirical 3-domain model in which the sensor, intervening HAMP, and catalytic domains can adopt kinase-promoting or inhibiting conformations, that are in allosteric communication. The catalytic and sensor domains intrinsically favor a constitutively ‘kinase-on’ conformation, while the HAMP favors the ‘off’ state; when coupled, they create a bistable system responsive to physiological [Mg^2+^]. Mutants alter signaling by locally modulating these intrinsic equilibrium constants and couplings. Our model suggests signals transmit via interdomain allostery rather than propagation of a single concerted conformational change, explaining the diversity of signaling structural transitions observed in individual HK domains.

## Introduction

Two component system sensor Histidine Kinases (HKs) are a conserved signaling module in bacteria responsible for sensing a myriad of environmental stimuli and orchestrating transcriptional responses along with their cognate transcription factors (Response Regulator, RR) (1, 2). These sensors are generally implicated in environment sensing, and are involved in multi-drug resistance (3–5) and as master regulators of virulence programing in pathogenic bacteria (6, 7). HKs are constitutive homodimers, which transmit signals through a series of intermediary domains to a cytoplasmic catalytic domain. While the lack of a full-length HK structure has hampered our understanding of the mechanism of signal transduction in these proteins, cytoplasmic domain structures have shed light particularly on the enzymatic core of this class of kinases. Several crystallographic snapshots of the autokinase domains of multiple HKs in various conformations (8–14), particularly CpxA, DesK, and VicK, have shown distinct conformations involved in autophosphorylation, phosphotransfer and dephosphorylation that may be conserved across this family. While these structures offer a conserved view of the catalytic cycle of the cytosolic autokinase domain (15), the question of how these proteins couple a sensory event on the other side of the membrane, and many nanometers away to the modulation of the activity of this domain remains unanswered.

This question is especially perplexing in light of the various modular architectures of HKs, involving the insertion of one or more signal transduction domains between sensors and the conserved autokinase domain. It is abundantly clear that the same conserved autokinase domain that defines this protein class can be regulated by a myriad of structural inputs, ranging from short alpha-helical dimeric coiled coils, to well-folded tertiary folds such as HAMP, PAS and GAF domains (16, 17) (**Figure 1**). Moreover, it is clear from the representation of these folds in diverse protein classes that these domains evolved independently of HKs and were co-opted pervasively into functioning HK architectures. Therefore, they are likely to serve a generalizable function that is robust to evolutionary selection, and the construction of physiologically relevant sensors optimally positioned to respond to environmental changes. While some intervening transduction domains have clearly annotated functions, such as the binding of intracellular ligands which are integrated into the sensory function of the HK, the requirement for other signal transduction domains remains obscure.

**Figure 1.**
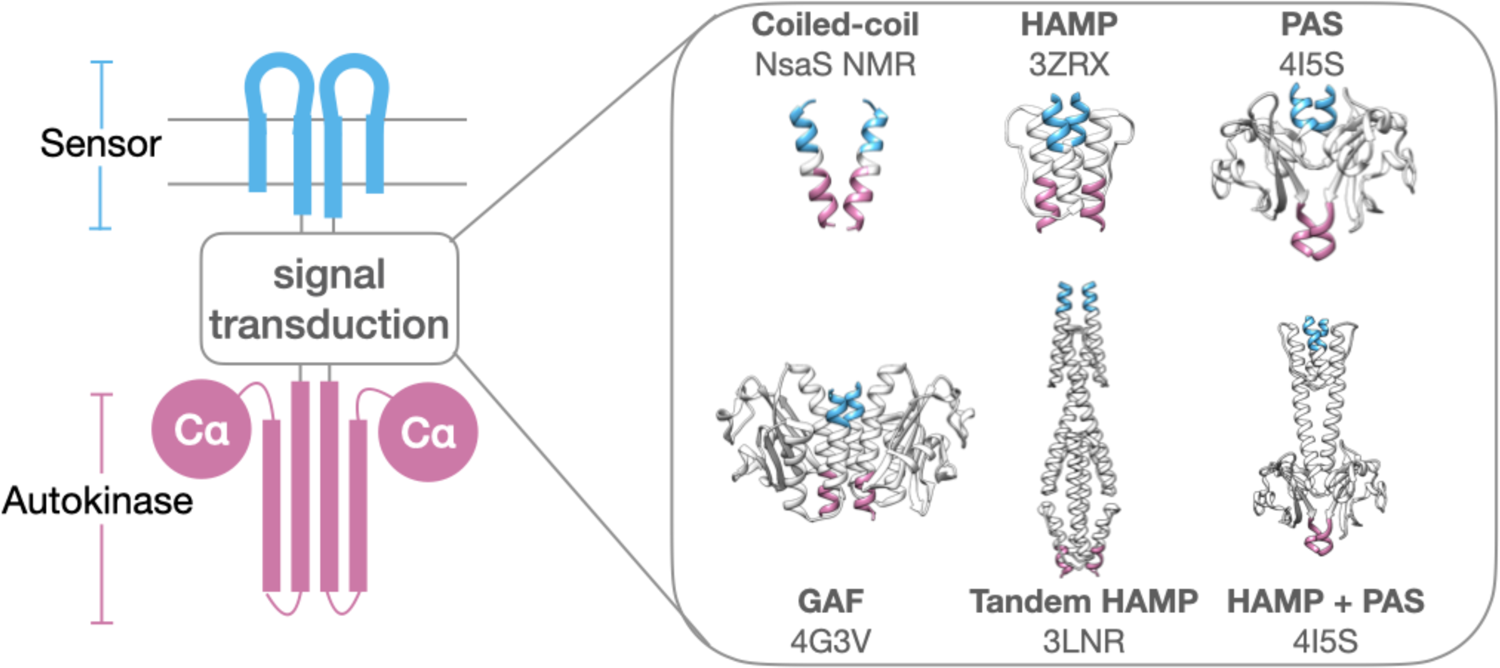
Modular architecture of histidine kinases. Various protein folds and numbers of signal transduction domains are found inserted between sensor (blue) and autokinase (purple). Structurally elucidated examples include simple coiled-coils (NsaS), HAMP (AF1503), PAS (VicK), GAF (Nlh2), Tandem HAMP (Aer2) and HAMP/PAS domain (VicK). Pdb codes are provided in figure, except for NsaS (NMR structure).

In this work, we evaluate the coupling of sensor and autokinase domains in a model Gram-negative HK, PhoQ (18), in which these domains are separated by intervening transmembrane and HAMP signal transduction domains. The PhoQP two-component system is composed of a canonical transmembrane sensor HK, PhoQ, that senses the presence of divalent cations (19, 20) and polycationic species such as antimicrobial peptides (21, 22), and a cognate response regulator, PhoP (18, 23), which transcriptionally controls regulons pertinent to cation transport and outer-membrane remodeling (24–33). The kinase activity of PhoQ is repressed by divalent cation binding, whereas it is enhanced by the presence of antimicrobial peptides. PhoQ is additionally implicated in low pH sensing (34) via an interaction with the membrane protein UgtL (35), and has more recently been suggested to respond to changes in osmolarity (36), although the mechanism is unclear. With respect to its most well characterized function, i.e., the sensing of divalent cations such as Mg^2+^, it is hypothesized that in the absence of such cations, the electrostatic repulsion between an acidic patch in the sensor domain and the negatively charged bacterial inner membrane enforces the ‘kinase-on’ conformation of the sensor and results in high-kinase/ low-phosphatase activity in the autokinase domain. In the presence of divalent cations, the electrostatic interaction between the sensor and inner-membrane are bridged, resulting in a different ‘kinase-off’ sensor conformation that corresponds to low-kinase/ high-phosphatase autokinase function (37, 38).

To probe the coupling between the sensor and autokinase domains, we established two assays, which allow simultaneous measurement of the conformational states of the sensor and autokinase domains (**Figure 2A**). Like most HKs, PhoQ is a constitutive parallel homodimer, in which the individual domains interact along a series of coaxial helical bundles. Previously, we observed that a Tyr60 à Cys variant forms interchain disulfides between the two monomers only in the absence of Mg^2+^ where the protein is in the ‘kinase-on’ state. Thus, the fraction of the sensor in the ‘kinase-on’ versus ‘kinase-off’ state can be readily quantified based on the amount of dimer versus monomer seen in a western blot. Importantly, the Y60C substitution is minimally perturbing, as the [Mg^2+^] dependent signaling curve for this mutant is nearly identical to wild type PhoQ with respect to the midpoint of the transition and activity of the basal and activated states. Also, the redox environment of the periplasm of *E. coli* is buffered such that disulfide formation is reversible and hence a good readout of the conformational state of the sensor. To quantify the activity of the auto-kinase domain, we use a well-established beta-galactosidase gene-reporter assay that employs the PhoQ/PhoP-controlled promoter of the Mg^2+^ transporter MgtA. Although this assay is indirect, there is a reasonable correlation between promoter activity and PhoP phosphorylation (39). We note that similar assays have been extensively used by Falke, Haselbauer et al. (40) to probe signal transduction in chemosensors that are related to PhoQ.

**Figure 2.**
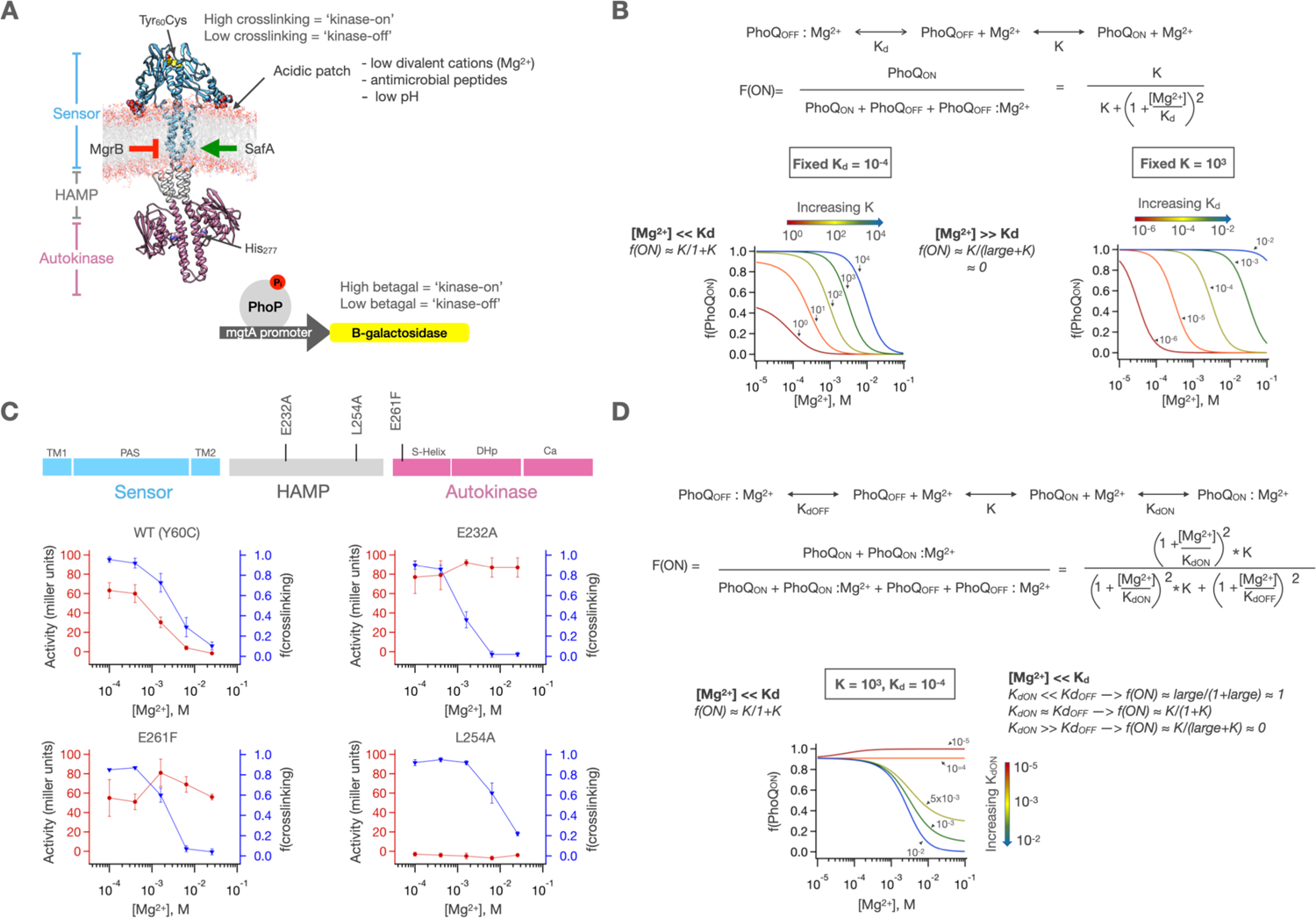
A fully concerted signaling mechanism does not explain PhoQ activity. A) MD model of PhoQ in which the sensor (res. 1-219, blue), HAMP (res. 220-260, grey) and autokinase domains (res 261-494, purple) are annotated. The sensor contains a Y60C mutation (spheres) which shows signal state dependent crosslinking. The autokinase contains the conserved catalytic His277, which upon phosphorylation transfers inorganic phosphate to the response regulator PhoP, which then modulates a *mgtA* promoter-driven beta-galactosidase reporter. Stimuli and regulatory proteins that modulate PhoQ activity are shown. **B)** A fully concerted (single domain) signaling scheme in which the intrinsic signaling equilibrium of PhoQ, K, is modulated by Mg^2+^ binding at affinity = K_d_. While this model allows for modulation of the low [Mg^2+^] asymptote and midpoint of transition, it does not allow for modulation of the high [Mg^2+^] asymptote. **C)** sensor crosslinking and autokinase activity determined for ‘wild type’ (Y60C) PhoQ (n=9), as well as 3 mutations (n=2) along the signal transduction pathway. The sensor state and autokinase activity are not identically correlating as would be predicted by a concerted signaling mechanism. Error bars correspond to ±SD error **D)** Allowing both states of PhoQ to bind Mg^2+^ allows for modulation of all 3 features, but still maintains obligatory correlation between sensor and autokinase.

Using this approach, we evaluate the extent to which the sensor’s conformational state couples to and dictates the conformational activity of the autokinase domain for a set of over 30 mutants, representing substitutions throughout the signal transduction pathway from the sensor to the autokinase domain. We show how these mutants can modulate the three basic characteristics of a PhoQ signaling response which need to fit the biological role of the HK- the signal strength at the limiting high and low [Mg^2+^], and the midpoint of the [Mg^2+^] dependent transition-over the physiologically relevant concentration ranges that *E. coli* encounters (0.1-10 mM). We further evaluate the intrinsic signaling equilibria of the sensor and autokinase domains by disrupting the allosteric coupling between them using poly-glycine insertions in the signal transduction pathway and show that both domains are highly biased to the ‘kinase-on’ state when uncoupled from each other. The intervening HAMP domain serves as a negative allosteric modulator of both these domains balances the stability of the ‘kinase-on’ and ‘kinase-off’ states, so that they can become responsive to physiological concentrations of Mg^2+^. With these concepts in mind, we establish, fit and evaluate a semi-empirical 3-domain allosteric coupling model to account for the sensor-autokinase coupling and high/low asymptote and midpoint of transition behaviors of 35 distinct point-mutant and poly-glycine insertions, and highlight the advantages of inserted signal transduction domains in robustly modulating the signaling behavior of HKs.

## Results

In the following sections, we first show how amino acid substitutions alter the signaling of PhoQ to induce a variety of outputs (**Figure 2**). We focus primarily on Ala substitutions at regions expected to be on the interior of the protein, and hence likely to alter the relative energetics of the kinase-promoting versus states. We also examined the effects of Trp substitutions in the TM helix at positions expected to map to the headgroup region of the biolayer, as similar substitutions often induce changes in signaling (**41**). We avoid substituting residues in the entire autokinase domain or residues in the sensor that are involved in Mg^2+^ binding.

For each of the 35 variants, Mg^2+^-dependent dose-response curves were evaluated for both transcriptional activation and crosslinking and were found to exhibit a rich range of behavior. Representative curves are shown in **Figure 2C**, and the entire collection of curves are shown in sections below. We seek the simplest allosteric model that can explain the entire gamut of phenotypes, by evaluating a series of simple allosteric models of increasing complexity. In the next sections, we use theoretical curves to qualitatively show the limitations of the simplest models. Ultimately, we develop a 3-domain model with variable coupling between the domains, which is sufficient to fully explain the entire range of activities of all 35 mutants.

### Fully cooperative two-state models are unable to explain the gamut of activities of mutants

The simplest model for signaling in HKs is one in which the entire HK exists in a two-state equilibrium of ‘kinase-on’ and ‘kinase-off’ states which is then modulated by ligand binding (**Figure 2B**). In the case of PhoQ, Mg^2+^ binding stabilizes the ‘kinase-off’ state. As such, the fraction of PhoQ that is ‘kinase-on’ at any given [Mg^2+^] is equal to the fraction of PhoQ that is ligand-free. In such a model, the upper asymptote of activity at low [Mg^2+^] ([Mg^2+^] << K_d_) is governed by K, the two-state equilibrium for the kinase-on versus kinase-on conformations ( f(active) ≈ K/(1+K), see **Figure 2** for definition of symbols). In the models considered in this paper, we assume that Mg^2+^ binds to single sites in the sensor domains. It is possible that binding between the sites is cooperative or that more than one Mg^2+^ ions are bound per domain. However, given the fact that the transcriptional assay is an indirect readout of the ‘kinase-on’ state, and as such is not necessarily perfectly linear with respect to the fraction of activation, we are not able to differentiate between models that differ subtly in their dose-response curves. However, our data (see below) are able to rule out highly cooperative models in which many binding sites must be occupied with high cooperativity (as in the Asp receptor (42)) as this would result in a much sharper dose-response curve. In the simple two-state model, the lower asymptote, PhoQ will always be pushed to a fully ‘kinase-off’ state at high enough [Mg^2+^] ([Mg^2+^] >> K_d_) because the fraction of autokinase in a given signal state is equivalent to the fraction of sensor in the corresponding state. The midpoint of transition is dictated by the relative magnitudes of K, which reflects the relative preference for the ‘kinase-on’ vs. ‘kinase-off’ state, as well as K_d_, which is the affinity of Mg^2+^ for the ‘kinase-off’ state, as shown in **Figure 2B**. Since the model does not allow for any decoupling of sensor and autokinase domains, the fraction of sensor that is crosslinked is identical to that of the autokinase in the ‘kinase-on’ conformation.

We next examined a large set of sensor and autokinase activities of ‘wild type’ (Y60C) PhoQ at 5 different concentrations of Mg^2+^ to evaluate whether they collectively deviate from the behavior expected for the simple two-state model. Illustrative data in **Figure 2C** show it is possible to have low levels of kinase-activity at low [Mg^2+^] even though the sensor remains in a high-crosslinking kinase-on state (e.g. L254A). Similarly, some mutations retain high kinase-activity in the autokinase despite the sensor transitioning to a predominantly low-kinase (low-crosslinking) state (e.g. E232A, E261F). Finally, some mutants produce higher levels of kinase activity at low-Mg^2+^ than ‘wild type’ PhoQ, demonstrating that even at the low-[Mg^2+^] conditions in which the sensor is fully in the crosslinked ‘kinase-on’ state, there remains a significant fraction of the WT autokinase that remains in the ‘kinase-off’ state (e.g. E232A is more active than WT PhoQ at low [Mg^2+^]). Therefore, this fully concerted signaling model is insufficient to describe the full range of activities of PhoQ variants.

It is possible to account for the variable kinase activity at high [Mg^2+^] by considering a model in which allows Mg^2+^ to bind to both the ‘kinase-on’ and ‘kinase-off’ states with different affinities, analogous to Mg^2+^ itself behaving as a low-affinity inverse agonist of PhoQ (**Figure 2D**). Indeed, for the sensor of PhoQ, there is no reason to preclude ligand binding in either sensor state, since the same negatively charged surfaces are present in both states and can conceivably still bind Mg^2+^, albeit at a much lower affinity due to the lack of bridging interactions (43). In this scenario, the ‘kinase-off’ asymptote at high [Mg^2+^] is determined by the relative values of K_dON_ and K_dOFF_, as shown in **Figure 2D**. However, this model cannot explain the large differences in the shapes of the curves seen for crosslinking versus reporter gene expression for the same mutant, as in **Figure 2C**. Such behavior requires an additional parameter that describes the degree of coupling between the sensor and autokinase domains.

### Allosteric coupling between sensor and autokinase domains

A ligand-dependent sensor can be allosterically coupled to an autokinase domain with a tunable coupling strength to allow for the desired degree of communication between the sensor and the autokinase. In such a scheme, the sensor would be a ligand-binding domain with all the properties previously described for a fully concerted HK. The autokinase, in the absence of linkage to the sensor, would have a constant activity level based on its own intrinsic ‘kinase off’ to ‘kinase on’ equilibrium. The sensor is then connected to the autokinase in a manner that biases the intrinsic autokinase equilibrium differently depending on which signaling state the sensor is in. A ligand-dependent allosterically modulated HK results from such a coupling, so long as sensor ‘kinase-on’ and ‘kinase-off’ states of the sensor alter the autokinase equilibrium differently. To reduce the number of parameters needed to describe such a model, we can define the intrinsic equilibria of the sensor and autokinase when they are connected to a reference state (e.g., ‘kinase-off’) with equilibrium constants as shown in **Figure 3A**. K_Sen_ is the ‘intrinsic’ equilibrium of the sensor domain when connected to an autokinase in the ‘kinase-off’ state, and K_AK_ is the ‘intrinsic’ equilibrium of the autokinase domain when connected to the sensor in the ‘kinase-off’ state. When coupled to the ‘kinase-on’ state of either domain, K_Sen_ and K_AK_ are scaled by a new factor, ⍺. **Figure 3B-C** illustrate the effect of ⍺ on the Mg^2+^ dose-response curves. When ⍺ = 1, the two domains are fully uncoupled, and the binding of Mg^2+^ to the sensor is unable to affect the autokinase domain (**Figure 3B**). A value of ⍺>1 means that when either of the domains switches to the ‘kinase-on’ state, the other domain’s propensity to switch ‘kinase-on’ state is also enhanced by that factor, creating a correlated ligand-mediated transition between sensor and autokinase (**Figure 3C**). If 0<⍺<1, then a transition to ‘kinase-on’ state is actually easier when the other domain is in the ‘kinase-off’ state, creating an anticorrelated ligand dependent behavior. When the absolute value of the log of ⍺ becomes very large (i.e., when ⍺ is either >>1 or approaching zero), the two domains are highly coupled (**Figure 3D**) and the system behaves as in the fully concerted 2-state models in **Figure 2**. Therefore, ⍺ is the coupling strength between the ‘kinase-on’ states relative to the coupling between the ‘kinase-off’ states built into K_Sen_ and K_AK_.

**Figure 3.**
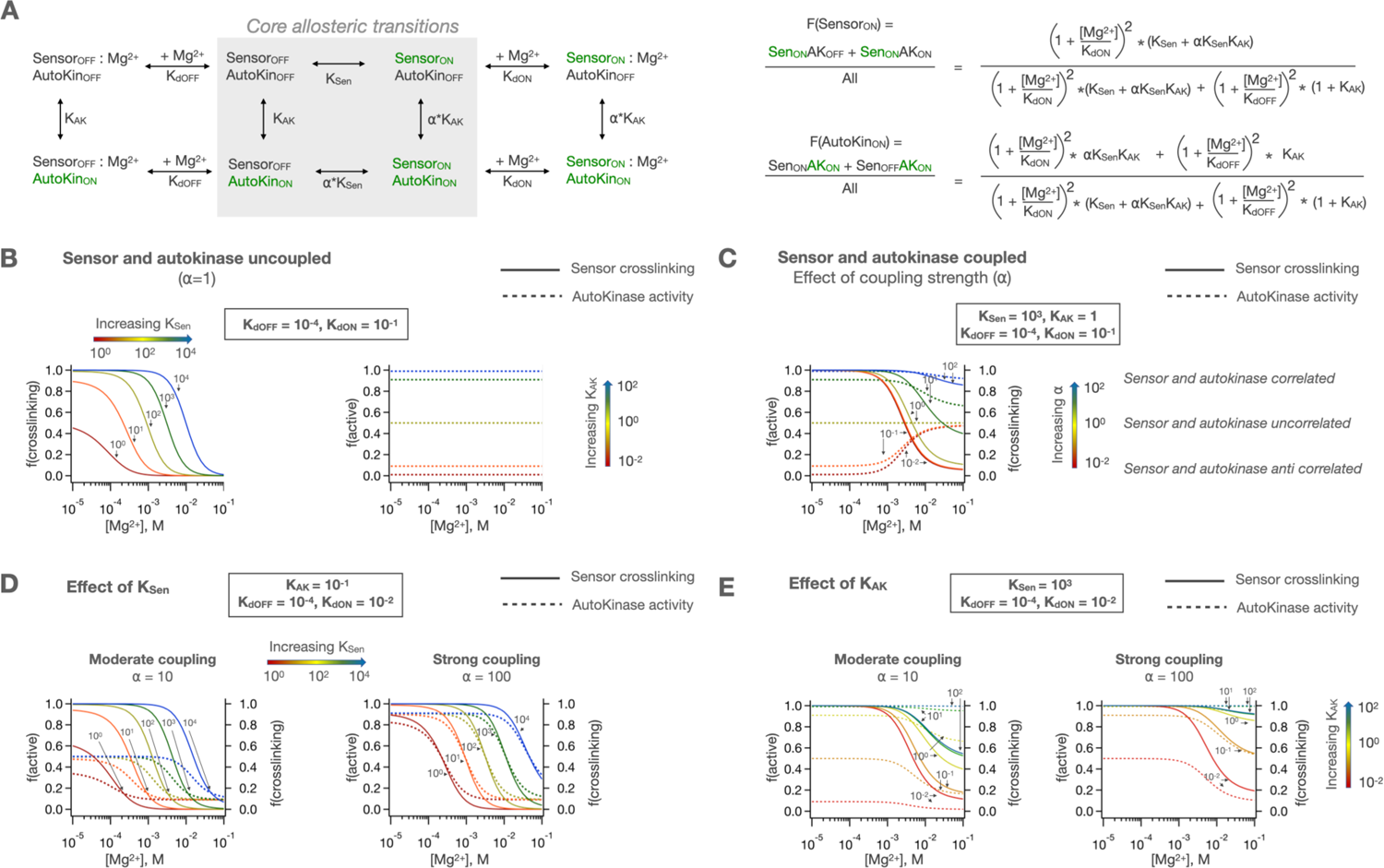
A 2 domain coupling scheme for PhoQ signaling. A) The sensor and autokinase domains of PhoQ are allowed to sample both ‘kinase-on’ and ‘kinase-off’ states with equilibrium constants K_Sen_ and K_AK_ when the other domain is in the ‘kinase-off’ state. When the other domain is in the ‘kinase-on’ state, the equilibria are scaled by the coupling constant, ⍺. This allows for semi-independent fractions of sensor and autokinase in the ‘kinase-on’ state, which are computed as shown. **B)** In the uncoupled case (⍺=1), K_Sen_ modulates the sensor identically to the previously described concerted signaling mechanism, while K_AK_ sets the basal autokinase activity. **C)** The coupling of these domains with ⍺≠1 results in [Mg^2+^] dependent activity that is either correlated (⍺>1) or anticorrelated (⍺<1). As ⍺ gets larger, the two domains act more as one concerted protein. **D)** Changes in the intrinsic equilibrium of the sensor affect autokinase activity through coupling, and similarly **E)** changes in the intrinsic equilibrium of the autokinase domain can alter the ligand-dependent crosslinking behavior of the sensor.

Coupling provides a robust mechanism for setting both the upper and lower activity asymptotes of the full-length sensor kinase. At high enough [Mg^2+^], the low-crosslinking ‘kinase-off’ state of the sensor becomes dominant, and the corresponding activity of the autokinase will be dictated by the autokinase equilibrium when coupled to this ‘kinase-off’ state, K_AK_. At low [Mg^2+^], the high-crosslinking ‘kinase-on’ state of the sensor will be dominant, and the corresponding activity of the autokinase will be dictated by ⍺*K_AK_. The midpoint of transition will depend on the relative magnitudes of all the parameters. However, the range of behaviors possible by this model of coupling depends heavily on the intrinsic equilibria of the sensor and autokinase themselves (K_Sen_, K_AK_). We next purposefully eliminated coupling experimentally (⍺=1) to obtain estimates of K_Sen_ and K_AK_, and to evaluate whether the 2-domain allosteric coupling mechanism is feasible in PhoQ.

### The effect of decoupling the HAMP domain from the catalytic and sensor domains

The allosteric model in **Figure 3** predicts that when the sensor and autokinase domains are fully uncoupled (⍺=1) these two domains act independently, i.e., the intrinsic equilibria of the sensor (K_Sen_) is the same irrespective of the state of the autokinase, and the equilibria of the autokinase (K_AK_) likewise becomes independent of the state of the sensor. To accomplish this decoupling experimentally, we insert a stretch of 7 helix-disrupting glycines (Gly_7_) to interrupt the helical connections that are required for coupling between PhoQ’s domains. One Gly_7_ insertions was introduced just before the HAMP (Gly_7_ −219/220) as the TM helix exits the membrane. The other Gly_7_ insertion (Gly_7_ −259/260) was made just after the HAMP signal transduction domain within a short helical connection to the autokinase domain. By comparing the effects of these insertions, we can decipher the role of the intermediate HAMP domain in signal propagation.

As expected, both mutants decouple Mg^2+^ binding from kinase activity (**Figure 4**). However, they have markedly different effects on the sensor and catalytic domains when these activities are evaluated individually. When the HAMP is decoupled from the sensor by introducing the Gly_7_ insertion between the HAMP and sensor domains, the sensor is highly activated, and remains in the high-crosslinking state, even at concentrations of Mg^2+^ sufficient to switch WT to the kinase-off state. On the other hand, if the HAMP remains coupled to the sensor as in (Gly_7_ −259/260) it behaves normally, being efficiently crosslinked in a [Mg^2+^]-dependent manner similar to WT. Thus, the HAMP would appear to favor the ‘kinase-off’ state, serving to reset the energetics of the otherwise highly stable ‘kinase-on’ state of the sensor. The resulting coupling provides an energetic balance so the system can respond to Mg^2+^ over the physiological range. The HAMP has a similar influence on the catalytic domain. When the native connection between the HAMP and the catalytic domain is disrupted by Gly_7_ insertion, it is highly activated. By contrast, when the connection between the HAMP and catalytic domains is retained as in WT, the kinase activity is strongly downregulated.

**Figure 4.**
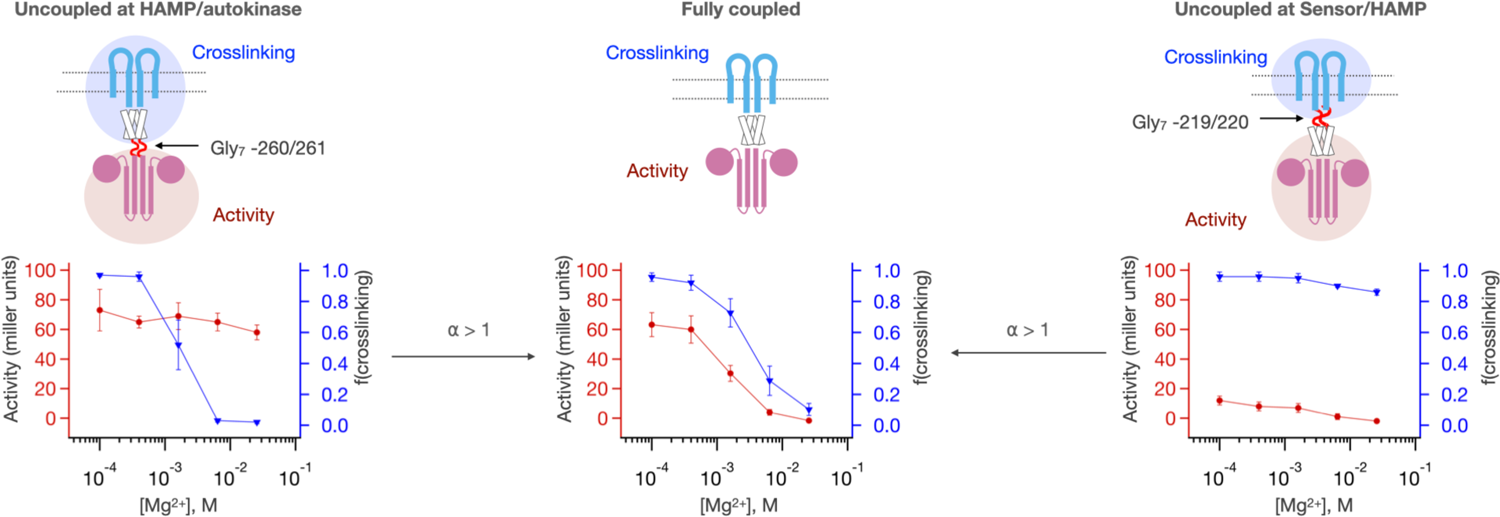
Intrinsic activities of the PhoQ sensor and autokinase domains are altered by coupling to HAMP. Gly_7_ disconnections are inserted either between the HAMP and the autokinase (Gly_7_ −260/261, left, n=3) or between the Sensor and HAMP (Gly_7_ −219/220, right, n=2) to disrupt allosteric coupling between sensor and autokinase. Both the sensor and autokinase by themselves have high ‘kinase-on’ propensity. However, presence of the HAMP with either of these domains potentiates the ‘kinase-off’ state, resulting in a more [Mg^2+^] responsive sensor, or a lower basal activity autokinase. These halves can be connected with positive allostery (⍺>1) to result in the observed sensor/autokinase activity of fully coupled PhoQ (middle, n=9).

These findings show that the HAMP domain serves not solely as a passive element that transmits a signal between the sensor and autokinase, but also plays a more active role in modulating the energetics of otherwise constitutively-on sensor and kinase domains so that the overall protein becomes fully responsive to physiological [Mg^2+^]. Thus, the HAMP domain serves as a separate, intervening domain with its own intrinsic equilibrium, in which the intrinsically preferred signaling state is negatively coupled to both sensor and autokinase.

We also see that the two-domain model in **Figure 3** captures much of the phenotypic behavior of the mutants. However, the fact that different effects are seen for decoupling before and after the HAMP indicates that it needs to be treated as a separate domain with its own equilibrium constant and coupling to both the sensor and catalytic domains. The treatment of the HAMP as a separate domain is of course parsimonious with its separate evolution as a bistable domain independent of histidine kinases, and its modular insertion into the various classes of proteins for which it is named. In subsequent sections we will additionally see this parsing into variably coupled domains can explain how mutations in the HAMP can modulate the sensitivity to Mg^2+^ and the dynamic range of PhoQ signaling.

### The HAMP domain is negatively coupled to the autokinase domains of CpxA and BaeS

Given the profound effect of the HAMP domain on the intrinsic activities of the PhoQ sensor and autokinase domains, we sought to examine if HAMP domains have similar effects in closely related but functionally distinct HKs with the same arrangement of signaling domains as in PhoQ. While we do not have facile means for evaluating the effect of the HAMP on sensor domains, we can examine the transcriptional activity of autokinase domains with and without coupling to their HAMP domains. We constructed Gly_7_ insertions in two closely related *E. coli* HKs, CpxA and BaeS, that have very similar architectures to PhoQ (both have similarly arranged PAS sensor, antiparallel 4-helix TM, a single cytosolic HAMP, and the conserved autokinase domains). The HK CpxA responds to periplasmic protein misfolding stress via an accessory protein, CpxP, and upregulates genes to mitigate this stress, such as periplasmic proteases and chaperones and modulation of outer membrane porin expression (44–46). It is similar to PhoQ in that the free HK is kinase-active, and is turned off by the binding of the periplasmic CpxP protein (47). BaeS is a closely related HK, which has significant overlap with CpxA, both in the inducing stimuli as well as the genes regulated (48). We evaluated the activity of these kinases using previously validated fluorescent gene-reporters (p*cpxP*::GFP for CpxA activity(49), p*spy*::mCherry for BaeS activity(50)) in a double CpxA/BaeS knockout strain. When the Gly_7_ motif is inserted immediately upstream of the autokinase domain, we observe a high basal activity for both kinases similar to PhoQ (**Figure 5**). However, when the Gly_7_ motif is migrated upstream of the HAMP thereby allowing the HAMP to couple to the autokinase, this high basal activity is potently repressed, again similar to PhoQ. This suggests that the HAMP strongly coupling to and altering the intrinsic activities of adjacent domains may be a generalizable principle, although it might not serve as a negative element in all cases.

**Figure 5.**
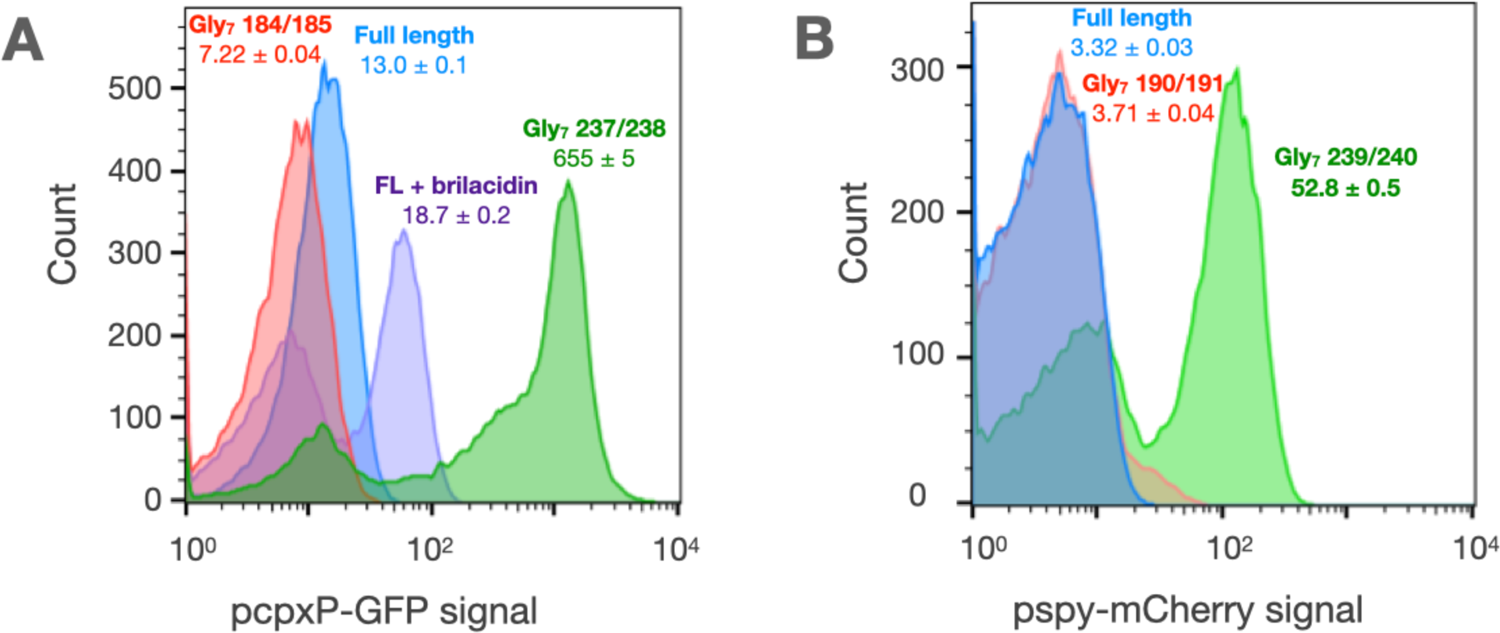
Glycine disconnections in CpxA and BaeS. A) The activity of CpxA constructs is measured in AFS51 strain (Δ*cpxA*) using a p*cpxP*::GFP reporter. Wild type CpxA is responsive to the antimicrobial mimetic, brilacidin (purple histogram). The autokinase domain of CpxA in isolation shows very high kinase activity (green), which is repressed to basal levels by the addition of the HAMP domain alone (red). **B)** The activity of BaeS constructs is measured in a Δ*baeS* Δ*cpxA* strain using a p*spy*::mCherry reporter. The autokinase domain of BaeS shows high kinase activity (green), which is repressed by the addition of the HAMP domain alone (red). Median reporter fluorescence values ± STE (n=20,000) are reported below labels for single experiment.

### Three-domain allosteric coupling mechanism of signal transduction

We next turned our attention to building a quantitative model that describes all the experimental data for 35 mutants. Based on the results of Gly_7_ insertion mutants, we developed a three-domain model, with the boundaries defined before and after the HAMP. In this model, the HAMP has its own intrinsic equilibrium, K_HAMP_, and there are two coupling constants that describe how the sensor couples to the HAMP (⍺_1_), and how the autokinase couples to the HAMP (⍺_2_). All possible state transitions are enumerated in **Figure 6A**. This treatment allows for semi-independent modulation of the sensor and autokinase using the intrinsic equilibrium of the HAMP. In the case where ⍺_2_ = 1, the autokinase is decoupled from the sensor + HAMP. In this scenario, the HAMP can modulate the [Mg^2+^] dependent state transition of the sensor through coupling via ⍺_1_ without altering the basal autokinase activity, as shown in **Figure 6B**. In the case where ⍺_1_ = 1, the sensor is decoupled from the HAMP+autokinase, and the HAMP can modulate the basal (and ligand-insensitive) activity of the autokinase through coupling via ⍺_2_, as shown in **Figure 6B**. When the protein is fully coupled (i.e. ⍺_1_, ⍺_2_ ≠ 1), we can potentiate the ‘kinase-on’ or ‘kinase-off’ states of the sensor and autokinase in a manner that depends on both K_HAMP_ and ⍺_n_’s, as shown in **Figure 6C**. Of particular interest is the case where ⍺_1_, ⍺_2_ <1, which enables the simultaneous potentiation of the ‘kinase-off’ state, while maintaining correlated sensor-autokinase behavior as observed in our Gly_7_ insertion experiments. Other possible behaviors with this 3-domain model include correlated sensing with ‘kinase-on’ potentiation, and anticorrelated signaling.

**Figure 6.**
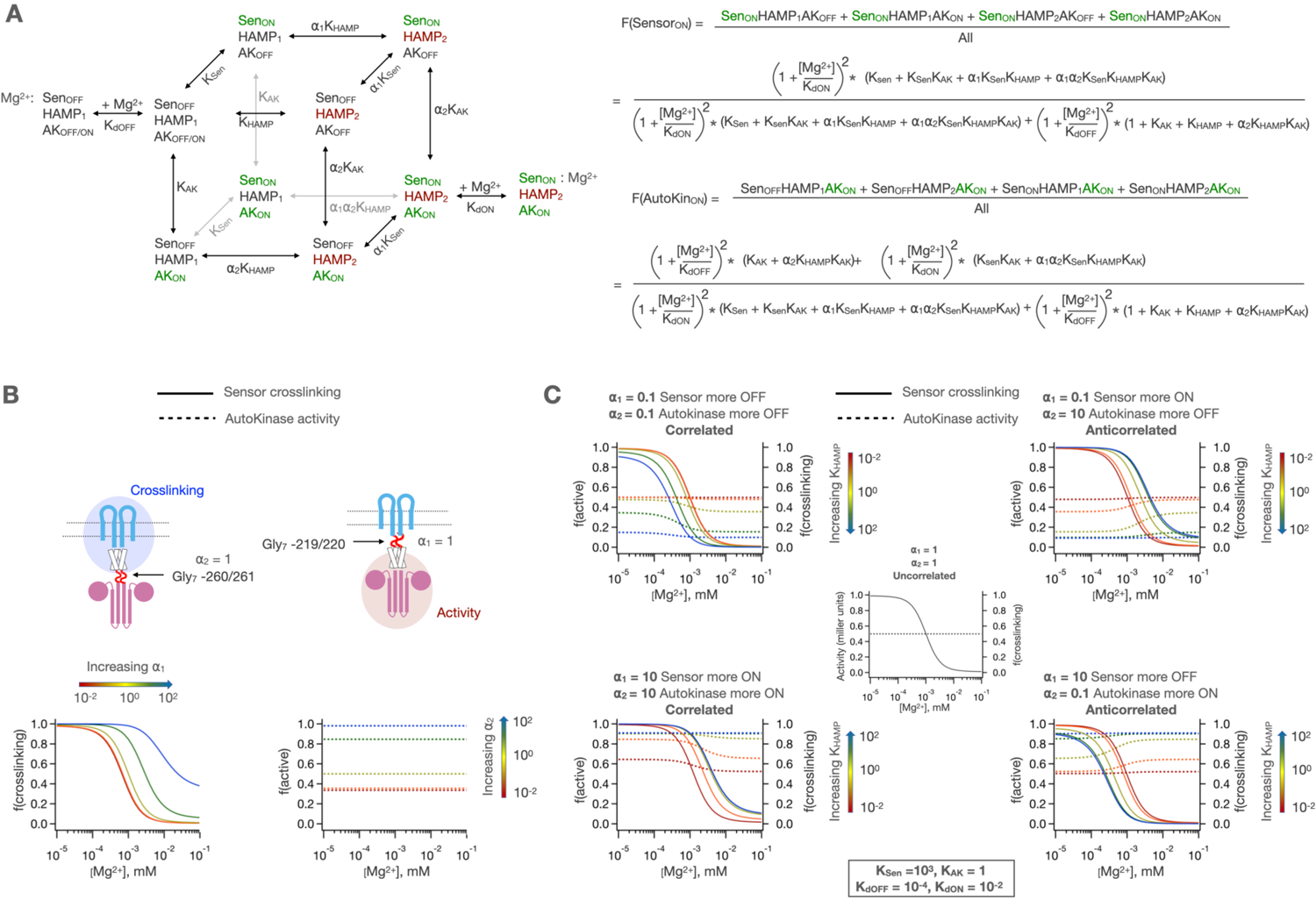
3-domain allosteric coupling model for PhoQ signaling. A) The sensor, HAMP and autokinase domains of PhoQ are allowed to sample both ‘kinase-on’ and ‘kinase-off’ signaling states while coupled to ‘kinase-off’ states in adjacent domains with equilibria K_Sen_, K_HAMP_ and K_AK_ respectively. When adjacent states are in ‘kinase-on’ states, the equilibria for transition are scaled by ⍺_1_ (sensor-HAMP) or ⍺_2_ (HAMP-autokinase). Predicted fraction of sensor crosslinking or autokinase activity are computed as shown below. **B)** The HAMP allows for the independent modulation of the basal state of the sensor or autokinase. When ⍺_2_ = 1, the HAMP modulates the [Mg^2+^] dependent transition of the sensor, and when ⍺_1_ = 1, the HAMP modulates the basal activity level of the autokinase. **C)** the two allosteric coupling constants allow for both correlated and anticorrelated modulation of sensor and autokinase and allow for potentiation of both the ‘kinase-on’ and ‘kinase-off’ states.

In order to fit our semi-empirical models to experimental observations, we generated a set of 35 single-point mutants and Gly_7_ insertions and *simultaneously* determined the sensor-crosslinking and autokinase activity at 5 different concentrations of Mg^2+^. We sampled regions all along the signal-transduction pathway between the sensor and autokinase, including the dimeric interfacial helices of the sensor dimer, the 4-helix transmembrane domain, the HAMP, as well as the conserved S-Helix motif that couples the HAMP to the autokinase (**Figure 7A**).

**Figure 7.**
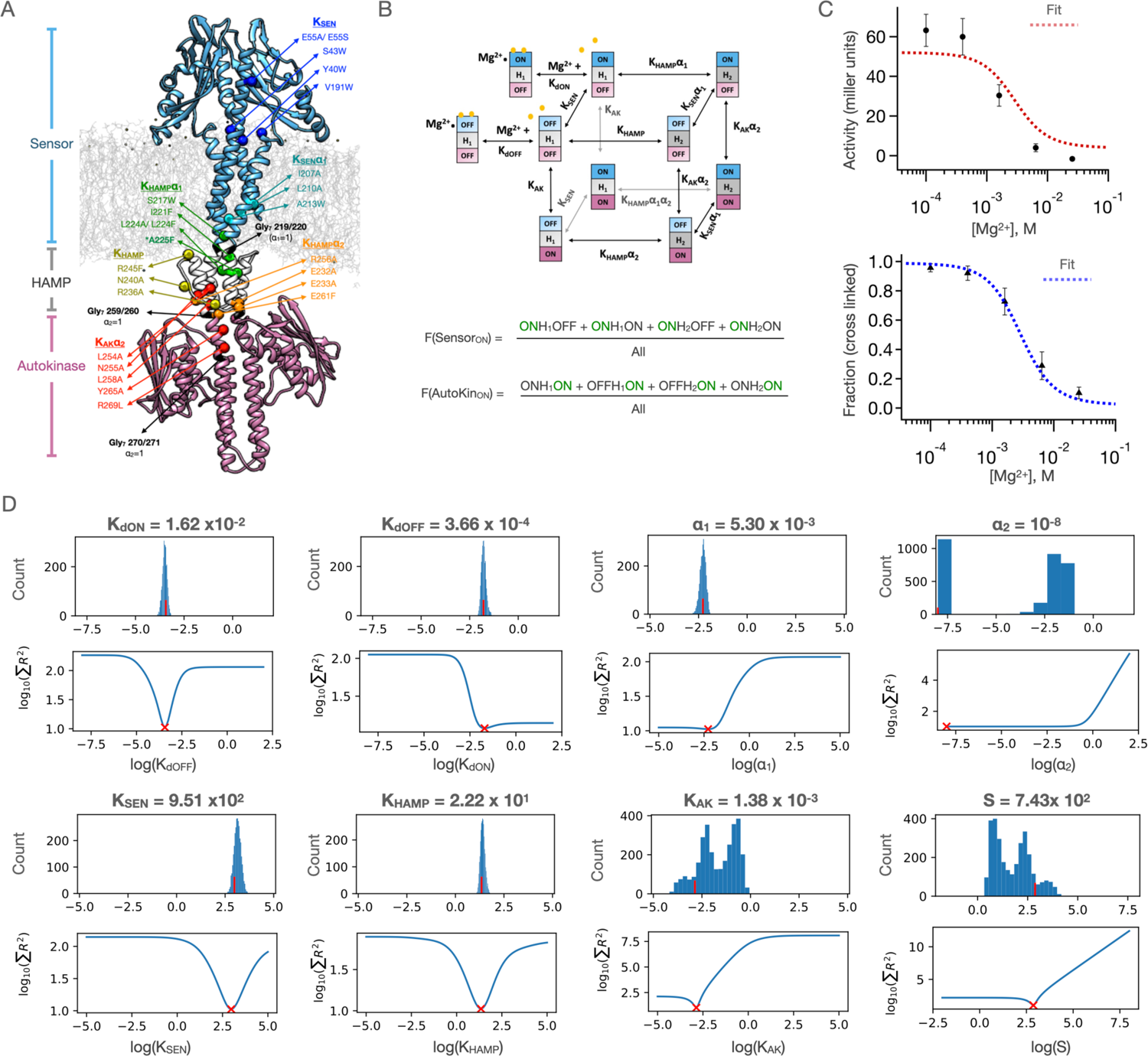
Results of 3 domain 2-state allosteric model fit of PhoQ activity. **A)** homology model of full length PhoQ showing positions of point mutations and Gly_7_ insertions used in study. Colored spheres represent the location of the C_β_ sidechain carbon, and are color coded by the identity of local parameters varied. **B)** 3-domain 2-state allosteric model used for fitting (see also Figure 6A) **C)** Fits to the [Mg^2+^] dependent activity (top) and sensor crosslinking (bottom) for ‘wild type’ Y60C PhoQ are shown. Error bars correspond to ± SD for n=9 biological replicates. **D)** Bootstrapped confidence intervals (top) and residual sweep analyses (bottom) are shown for all 8 global parameters. The value of the fit is indicated with red (x) and (|) marks. The confidence intervals of parameters S, K_AK_ and ⍺_2_ are further parsed in **Figure 7-figure supplement 1**.

**Figure 7 figure supplement 1.**
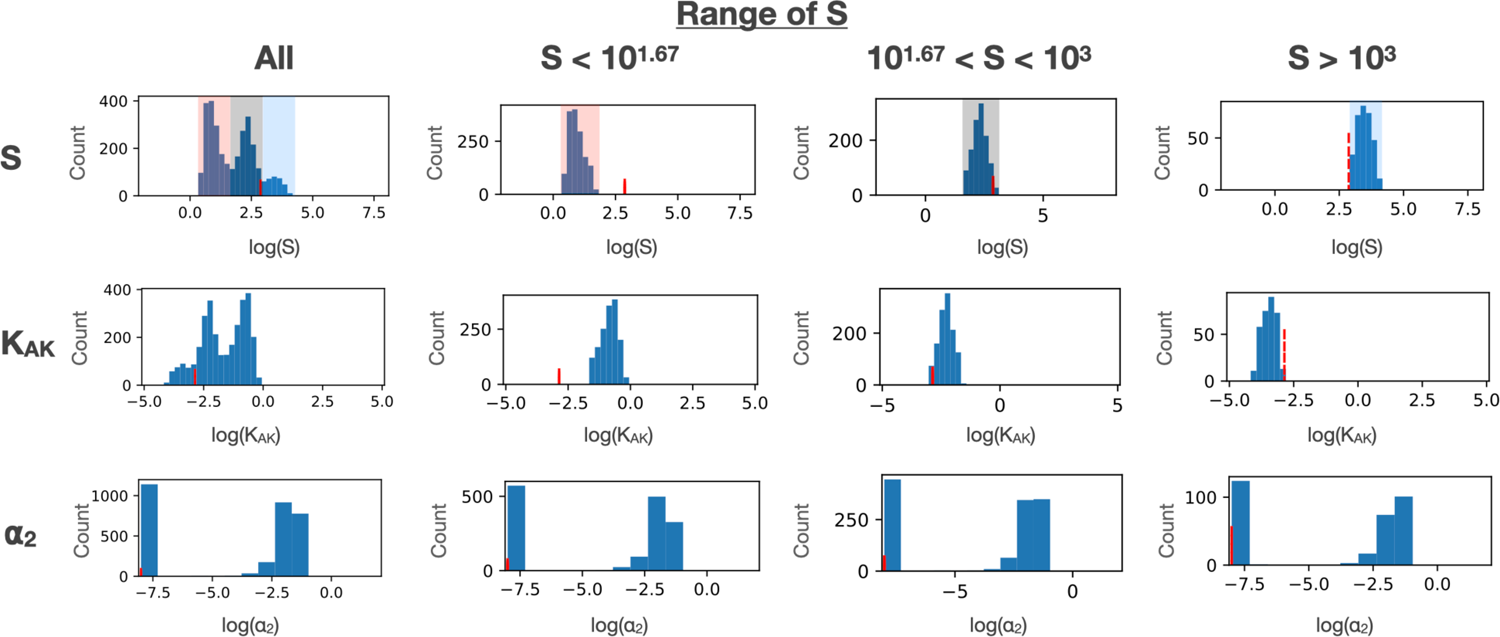
Effect of constraining S and K_AK_. The confidence intervals for K_AK_ and ⍺_2_ parameters are shown as a function of different ranges of S values. S and K_AK_ show strong correlation. **Application of the 3-domain model to a set of mutants illustrates how substitutions distant from active sites modulate signal strength and ligand sensitivity.**

Using this set of mutants, we next sought to determine the five core allosteric parameters (K_Sen_, K_HAMP_, K_AK_, ⍺_1_, ⍺_2_), and the dissociation constants for Mg^2+^ to the two sensor states (K_dOFF_, K_dON_). One last parameter (S) is a scaling factor that relates the mole fraction of autokinase in the ‘kinase-on’ state to the experimentally observed Miller units associated with the beta-galactosidase transcription (**Figure 6A**), which were obtained under strictly controlled experimental conditions to assure uniformity between mutants. In all, we sought to determine eight constants for each mutant. However, given the spacing of the points in our dose-response curves, it is only possible to obtain three pieces of information, i.e., the top, bottom and midpoint of the curves. Thus, with only six pieces of information (3 each from crosslinking and transcriptional activation) for each mutant, the model is under-determined for any one mutant.

We avoid this problem by using global fitting. For a given mutant, only one or two (or in 1 occasion, three) of the parameters are allowed to vary, with the others being fit as global parameters that are shared with other mutants. The choice of which parameters to vary is determined by the location of the perturbation on the primary sequence of PhoQ (**Figure 7A**, see methods). For example, a mutant near the N-terminus of the HAMP domain would be expected to primarily alter ⍺_1_ and K_HAMP_, so these values were allowed to vary locally. Mutants near the center of the tertiary structure of the HAMP domain are allowed to vary K_HAMP_ alone and so on. This results in an overall fit with 62 adjustable parameters corresponding to 8 global parameters, 47 locally varied parameters, and 7 parameters fixed to a value of 1 to account for Gly_7_ insertions (**Table 1**). By comparison, there are 6 * 36 = 216 observables. Thus, in theory, the data should be more than sufficient to define the independent parameters.

**Table 1.**
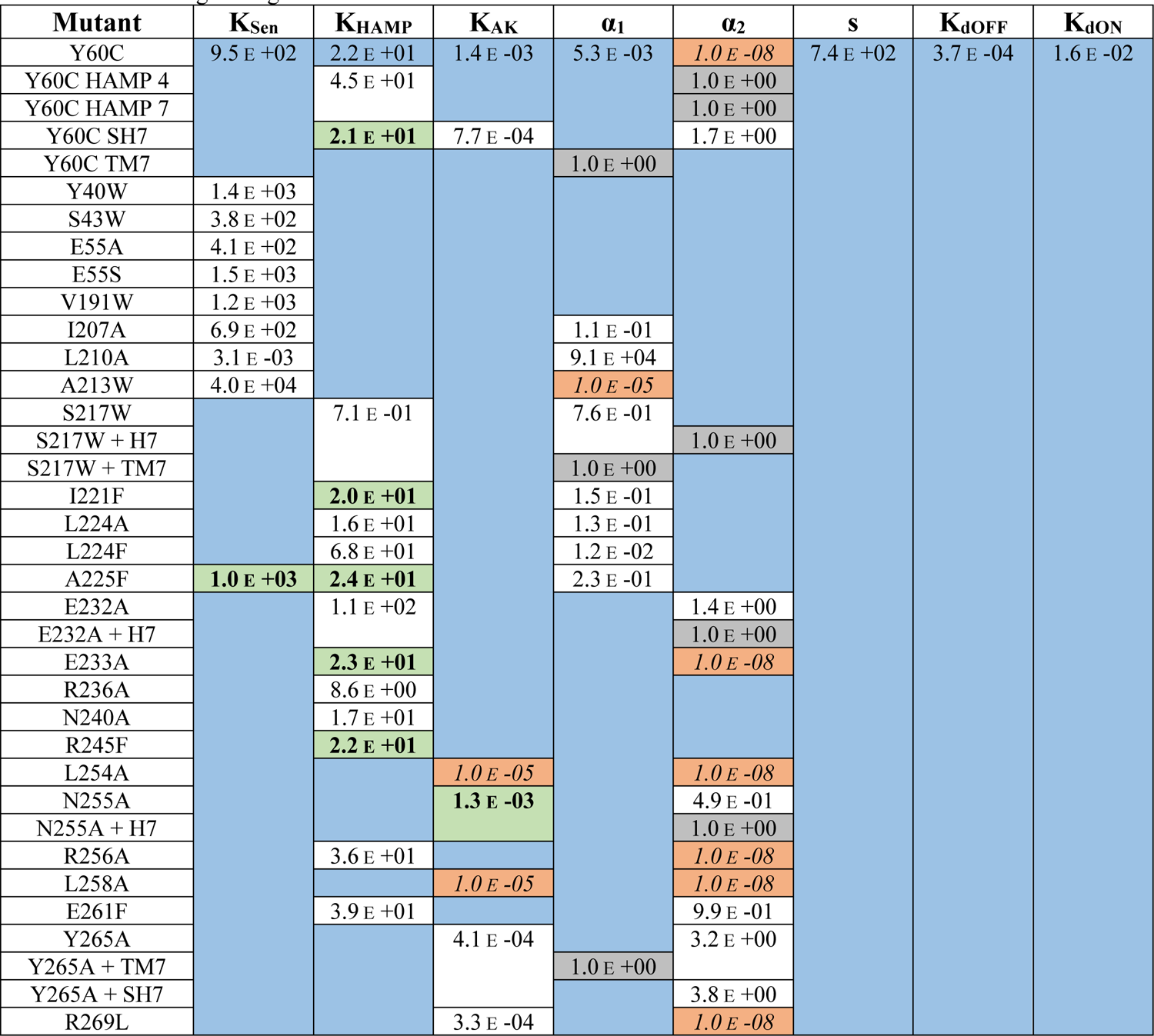
List of mutant parameter fits. Local parameters whose values remained within 10% of the global fit value are highlighted in bold font and green background. Parameters whose value drifted to one end of the explored parameter range are highlighted in italicized font and orange background.

This model was globally fit using our mutation dataset as explained in detail in the methods section. Briefly, we standardize the ranges of autokinase activity measurements (Miller units from beta-galactosidase assay) by the global average activity in our dataset. This normalizes the range of autokinase activity to one that is similar to crosslinking fractions (range 0 to 1) and gives both types of experimental measurements similar weights in our global fits. We give additional weight to data with experimental replicates (and hence greater certainty) by simply treating each replicate as an independent data set, with all the variables held constant between replicates during fit. Each parameter is allowed to sample a 10-log range of possible values, and the best fit is determined by minimizing the sum of residuals across the entire dataset. In order to avoid getting trapped in any local minima of the parameter space, we repeat the fit 125,000 times using randomly generated starting values for each parameter and determine confidence intervals for our parameters using a bootstrapping to generate over 3000 synthetic dataset fits (see methods for details). Where mutations or insertions have been introduced, we allow the parameters expected to be affected by the mutation to vary locally for the corresponding data set. Moreover, six mutants can be fit with fewer local parameters than were utilized in the fit, as the values for some of these locally fit parameters remain close to the globally fit value (within 10%), as highlighted in **Table 1** (green).

We are able to obtain a remarkably good fit for our entire dataset with the aforementioned considerations. **Figure 7C** shows the results of the best obtained fit for ‘wild type’ PhoQ (Y60C) sensor-crosslinking and autokinase activity. Since the wildtype data is fit entirely globally, it represents the most stringent test for the performance of our model overall, and qualitatively shows good agreement between model fit and experimental data. The values of the eight global parameters corresponding to this wild type fit are shown in **Figure 7D**, alongside two metrics of fit quality. The first metric is a bootstrapped confidence interval, with the frequency histogram of resulting fit values shown in the top panels. The second metric is a parameter sweep analysis in which the global sum of residuals is evaluated as the value of the indicated parameter is allowed to vary while all other parameters are held fixed. Five of our global parameters, K_dOFF_, K_dON_, K_Sen_, K_HAMP_ and ⍺_1_ show excellent convergence to the ‘best fit’ value, with well-defined minima in the sum of residuals as we explore parameter value. Three parameters, K_AK_, S and ⍺_2_ show strong signs of covariability, and wider confidence intervals. In the fully activated state, the observed signal is defined by the product of the scaling factor, S, and the fraction of the protein in the ‘kinase-on’ state (approximately S*K_AK_). This product is well-defined and converges to a value of ≈ 1.02. However, as K_AK_ is lowered below this value, S increases in parallel to maintain a constant value for the product of S*K_AK_. In **Figure 7-figure supplement 1**, we show that when the values of S are restrained, the values of K_AK_ are also restrained, and vice versa. Nevertheless, we can place a mechanistically meaningful upper limit on K_AK_, of approximately 0.1. Similarly, we can place an upper limit on the value of 0.1 for ⍺_2_, which represents the negatively cooperative coupling of K_AK_ to the parameters defining the other domains. These uncertainties do not affect any of our conclusions below, which depend on presence of strong versus weak and negative versus positive coupling.

One feature that was somewhat surprising was that K_AK_ is actually unfavorable towards forming the ‘kinase-on’ versus ‘kinase-off’ states (K_AK_ < 1), even at limiting low concentrations of Mg^2+^. This indicates that the observed activity for the WT protein is less than what is observed for some of the mutants, and what might be observed in a hypothetical state in which the autokinase is unfettered by connections to HAMP and the membrane. Although unexpected, this finding is consistent with a large body of data (51–53), and has been observed in PhoQ with antimicrobial peptide stimulation (54). Thus, in ligand-responsive HKs, evolution does not drive towards maximal activity which might lead to wasteful and toxic transcription, but instead a finely tuned value that is titrated to the degree of transcription required for function.

The values of the parameters provide a detailed view of the energy landscape of PhoQ, in the ‘kinase-on’ and ‘kinase-off’ state – and how it is modulated by binding to Mg^2+^ and mutations. The parameters are consistent with our earlier observations that the HAMP is a significant modulator of the intrinsic equilibria of the sensor and autokinase domains. At high Mg^2+^ concentrations, PhoQ is in a ‘sensor-off’ and ‘autokinase-off’ state. With respect to this reference ‘kinase-off’ state, the HAMP domain has a thermodynamically favored signaling state, ‘HAMP_2_’, with a fit equilibrium value of K_HAMP_ = 22. This favored state of the HAMP is more strongly coupled to these ‘kinase-off’ states and serves to dampen the otherwise favorable transitions of both the sensor and autokinase domains to the ‘kinase-on’ conformation. The sensor’s propensity to switch to a ‘sensor-on’ state is reduced from a highly preferred equilibrium K_Sen_ = 9.51 x 10^2^, to a modest downhill equilibrium of ⍺_1_K_Sen_ = 5.0 when the HAMP is in this HAMP_2_ state. This latter equilibrium is weak enough to be overcome by Mg^2+^ binding, and the ‘sensor-off’ state is further stabilized with more ligand binding. The ‘HAMP_2_’ state that is preferred in this state is also strongly coupled to the ‘kinase-off’ state of the autokinase, reducing the propensity of the autokinase to switch to the ‘kinase-on’ state from S.K_AK_ = 1.0 to ⍺_2_.S.K_AK_ ≤ 10^-3^. Thus, the HAMP_2_ state behaves as a negative modulator of the intrinsic propensities of the sensor and autokinase. At high enough [Mg^2+^], the entire population ensemble is predominantly in the sensor_OFF_-HAMP_2_-Autokinase_OFF_ state.

In the absence of ligand, the sensor’s modest downhill equilibrium to the ‘kinase-on’ state is strongly tied to a switch of the HAMP from ‘HAMP_2_’ to ‘HAMP_1_’, with an equilibrium = 1/(⍺_1_K_HAMP_) = 25. The HAMP_1_ state is weakly coupled to the autokinase, which allows the autokinase to sample both kinase-off and kinase-on state, with an effective equilibrium of S.K_AK_ = 1.02. This allows for the partial decoupling of the sensor and the autokinase at low Mg^2+^ concentrations observed in wild type PhoQ with the population ensemble composed of both sensor_ON_-HAMP_1_-Autokinase_ON_ and sensor_ON_-HAMP_1_-Autokianse_OFF_ states. Because the HAMP_1_ state is weakly coupled to both adjacent domains, the equilibria constants K_Sen_ and K_AK_ are close to the intrinsic equilibria of these domains when uncoupled form the HAMP altogether. In other words, K_Sen_ = 9.5 x10^2^ reflects the high propensity of the sensor to switch to the ‘sensor-on’ state when uncoupled from the HAMP, and S.K_AK_ = 1.0 reflects the propensity of the autokinase to have as high an activity as full length PhoQ at low [Mg^2+^], as shown earlier with Gly_7_ disconnections in **Figure 4**.

The parameters for individual mutants show how amino acid substitutions alter the energy landscape and how these changes in turn alter the phenotype. Before discussing the effects of substitutions, however, it is important to address the overall quality of the fit over the full ensemble of mutants. **Figure 8** shows the results of fits for our mutations and Gly_7_ insertions, and the locally varied parameters. The corresponding fit values are listed in **Table 1**, and confidence intervals and parameter sensitivity analyses are shown in **Figure 8-figure supplement 1**. We obtain fits within experimental error for the [Mg^2+^]-dependent transcriptional activity of our entire mutant data set. Thus, the model works well for all functional mutants. The only deviations lie in the crosslinking data for non-functional mutants that are decoupled in the transcriptional assay (**Figure 8-figure supplement 2**). One such set of mutants (I221F, L224A and A225F) have substitutions at the C-terminal end of the second TM helix. While the midpoint and lower limit were well described by the model, the experimentally observed extent of crosslinking reaches an upper limit of 65% to 80% crosslinking at low Mg^2+^, less than the predicted value near 100%. Given the location of the substitutions near the membrane interface, it is possible that a portion of the protein is not fully inserted and hence the samples used for western analysis might have been contaminated by cytoplasmically localized, and not yet membrane-inserted protein, which would be expected to remain un-crosslinked. There are also two mutants localized near the interface between the HAMP and the autokinase domains where the mid-point is poorly fit (L254A, L258A), potentially owing to our choices of parameters to locally float for these mutants (**Figure 8-figure supplement 2D-E**) as discussed in methods. Significantly better fits were obtained by altering the parameters varied for these mutants from K_HAMP_ and α_2_ to K_AK_ and α_2_. Thus, these residues may be involved in the underlying equilibrium of the autokinase domain itself due to their proximity to the “S-helix” that connects the HAMP to the autokinase domain. Finally, double mutants are not fit well, especially when the two sites of mutation are in close proximity (S217W + HAMP Gly_7_, N255A + HAMP Gly_7_, Y265A + Sensor Gly_7_, **Figure 8-figure supplement 2F-H**). This is likely because the thermodynamic effects of double mutants are often non-additive in structurally and sequentially proximal positions that interact directly. Additionally, while we observe relatively invariant expression of almost all variants (as seen in the western analysis used to quantify crosslinking), some variants, particularly double mutants required slight induction of expression with 10 µM IPTG for observable levels of membrane-inserted PhoQ by western-blotting (see methods). In summary, the crosslinking and transcriptional activity data are very well fit for the entire set up mutants, except for a fraction of the nonfunctional mutants in which Mg^2+^ binding and transcription were significantly decoupled. Even for these mutants, however, there is a qualitative fit to the data, and likely reasons for the deviation.

**Figure 8.**
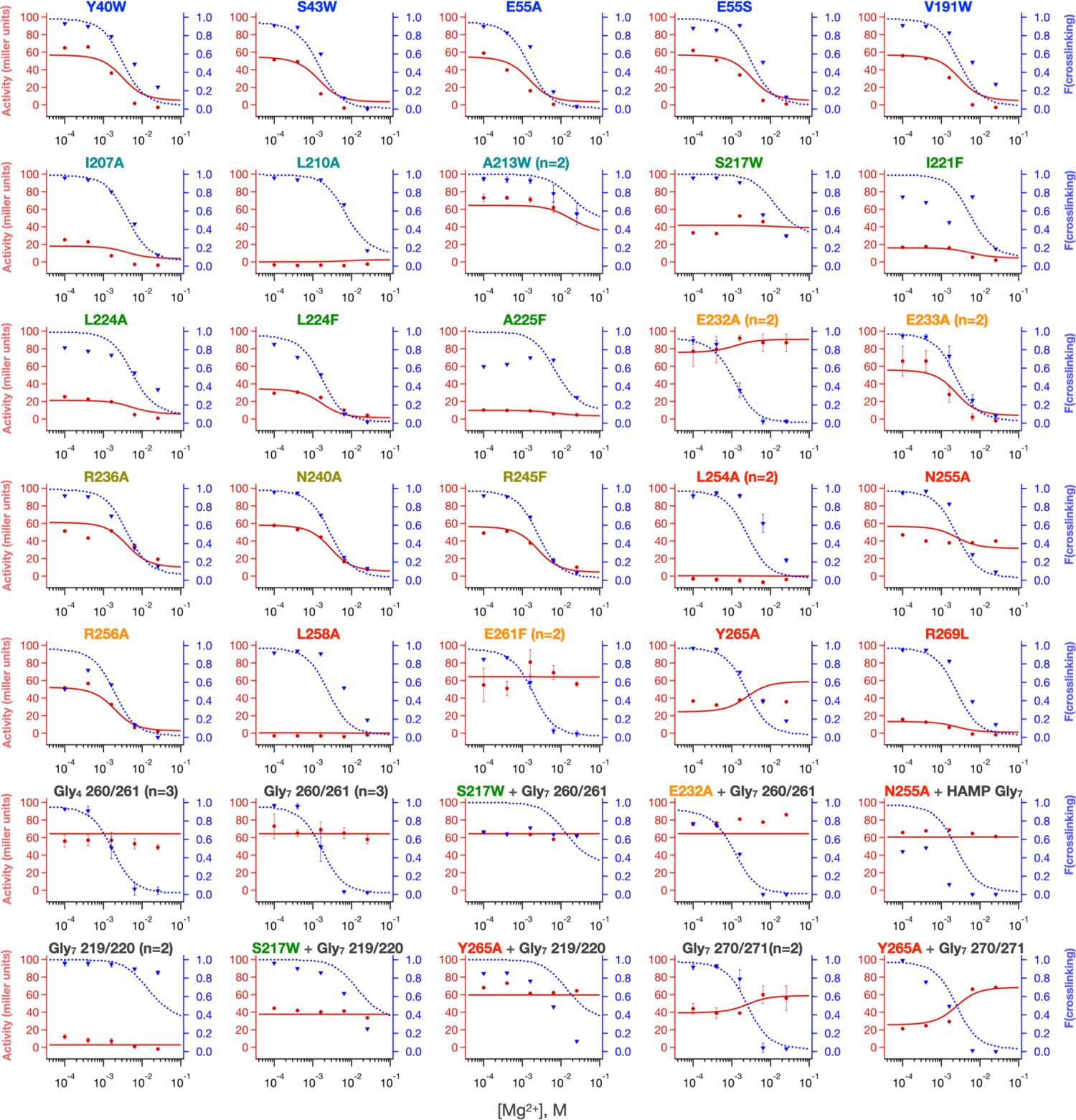
Local fits of sensor crosslinking and kinase activity for 35 PhoQ mutations. Fits to activity (red line, closed circles) and sensor crosslinking (blue dashed line, triangles) are shown for the entire PhoQ dataset. The identity of locally varied parameters is listed in Figure 7A and **Tables 1** and **2**. Confidence intervals and residual sweep analyses are presented in **Figure 8-figure supplement 1**. Poor fits are highlighted in **Figure 8-figure supplement 2**.

**Figure 8 figure supplement 1.**
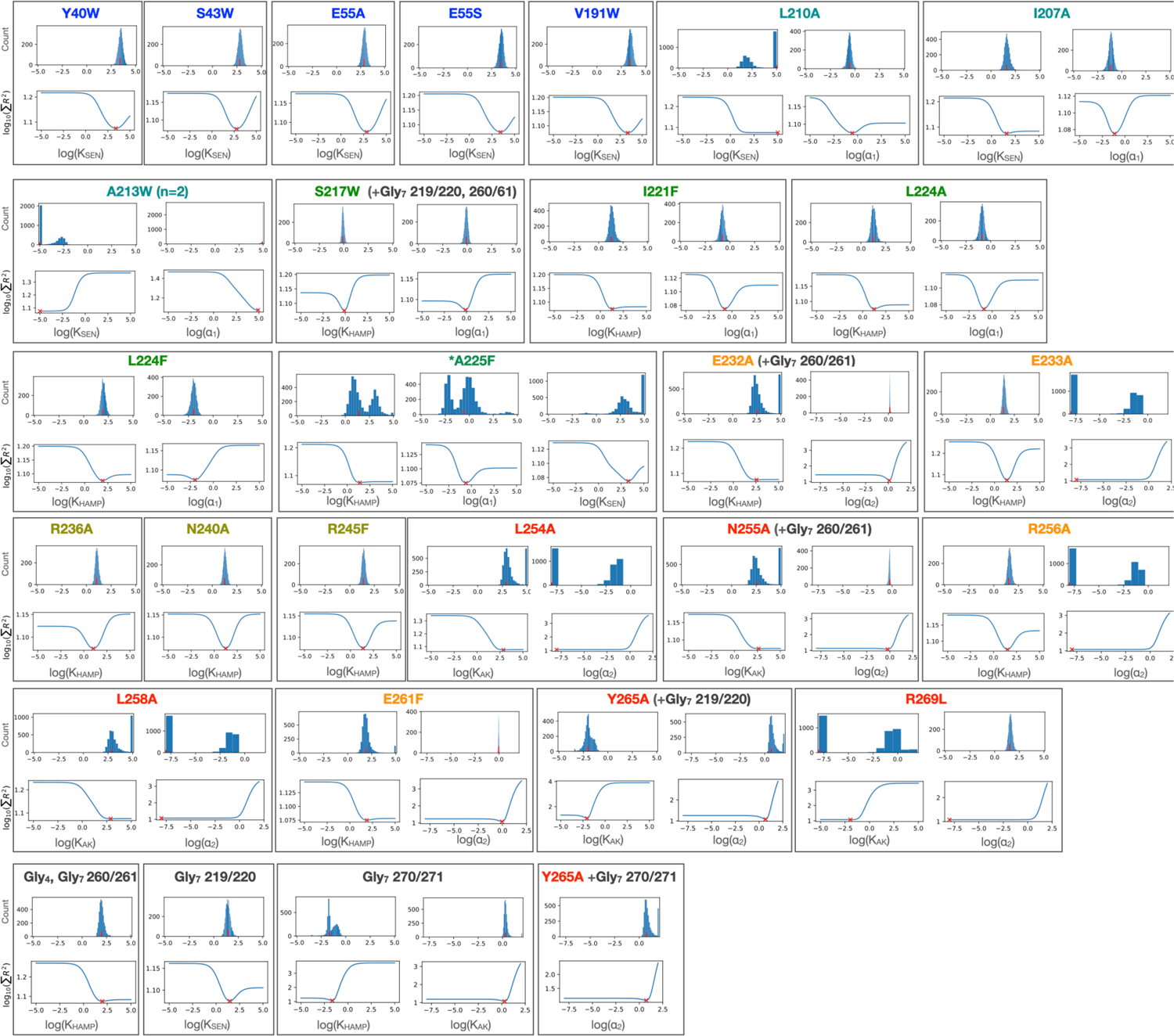
Bootstrapped confidence intervals and residual sweep analyses for PhoQ mutant fits. Histograms from 3061 convergent fits of simulated datasets for each local variable are shown in top panels. Residual sweeps in which the sum of residuals of the global fit is plotted as a function of indicated parameter being varied locally is shown in the bottom panels. Values of parameter fits are shown with red (x) and (|) marks.

**Figure 8 figure supplement 2.**
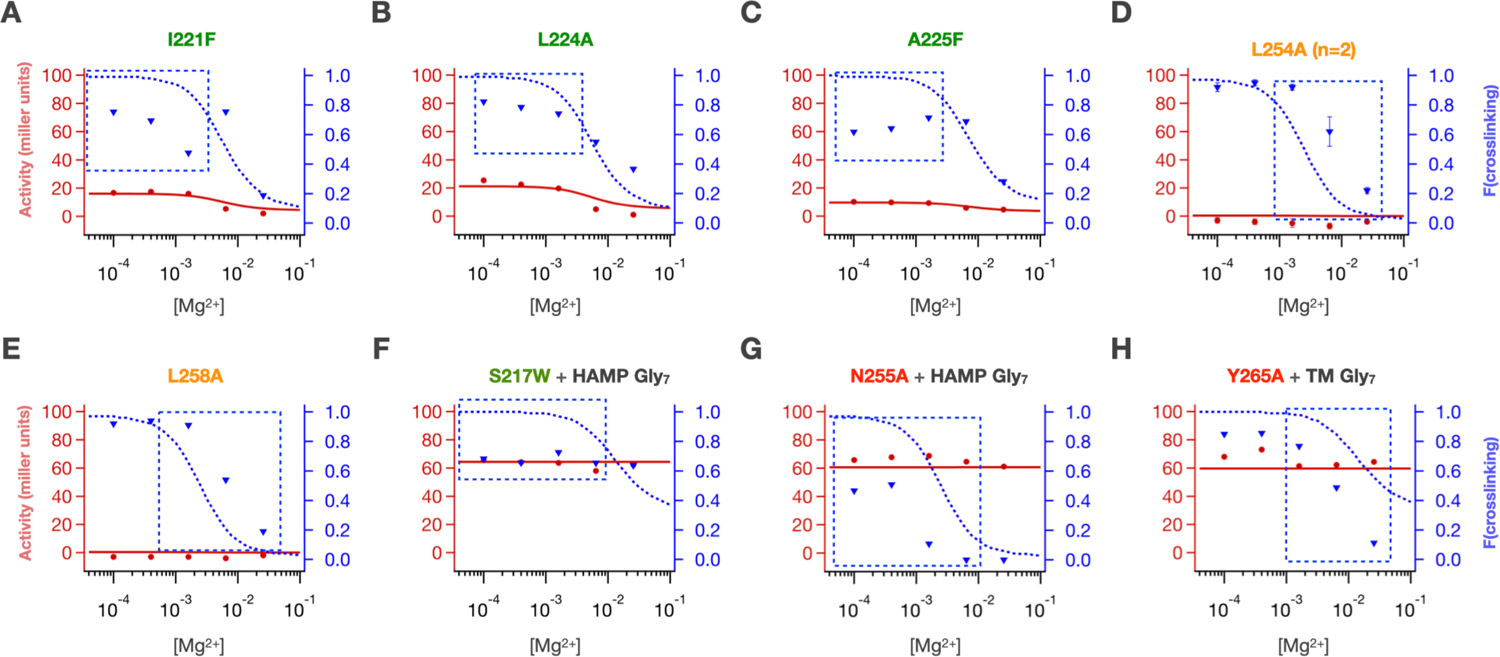
Poor fits were obtained for crosslinking at low [Mg^2+^] for **(A)** I221F, **(B)** L224A and **(C)** A225F. Poor midpoints of crosslinking transitions were fit for **(D)** L254A and **(E)** L258A. Some combinations of mutations had poor crosslinking fits **(F)** S217W + HAMP Gly_7_, **(G)** N255A + HAMP Gly_7_ and activity fits **(H)** Y265A + sensor Gly_7_

Our results illustrate the coupling of sequence and energetic landscapes in response to single-site substitutions. Although we chose a collection of mutants that were not involved in Mg^2+^ binding and catalysis, we observed a large range of effects on the transcriptional response of the mutants, including an inverse response in E232A and Gly_7_ 270/271 insertion. The advantage of the current analysis is that it shows how these mutations are able to alter the energetics of individual domains, and their coupling to adjacent domains. As an organism evolves to match its environment, its sensory systems need to adjust to the ligand-sensitivity (midpoint of the dose-response curve), the magnitude of the increase in the response (in this case, the (activity in the absence of Mg^2+^/activity in presence of Mg^2+^) and the basal activity (in the presence of saturating Mg^2+^). We consider these features separately.

**Figure 9.**
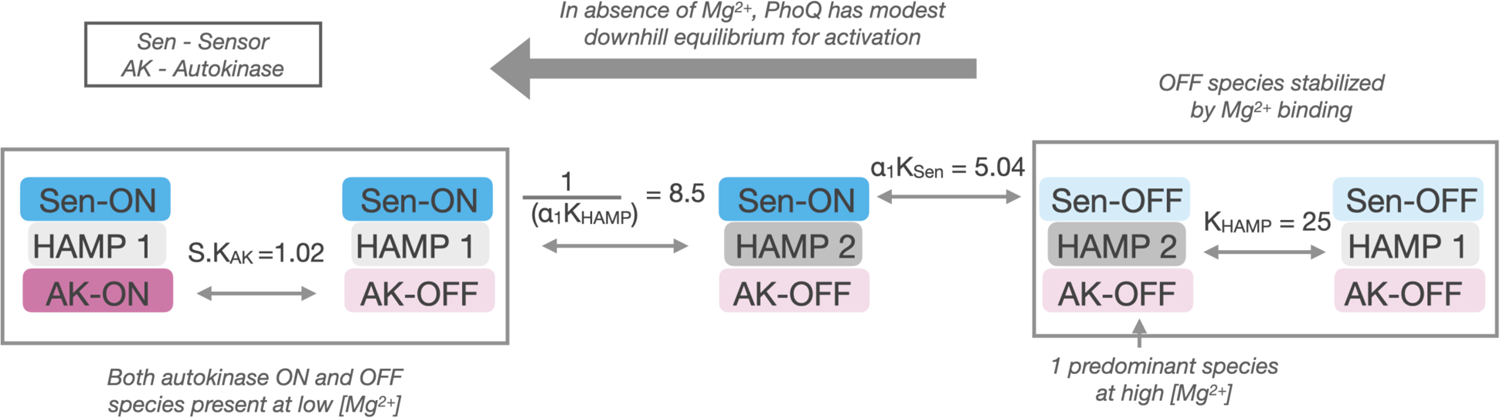
Allosteric pathway for PhoQ activation. In the absence of Mg^2+^, PhoQ has a moderate downhill equilibrium to a mixture of active states. Mg^2+^ binding is sufficient for overpowering this equilibrium and stabilizing the ‘kinase-off’ state, resulting in a predominantly Sensor-off/HAMP2/Autokinase-off population.

In a well-coupled system such as WT PhoQ, the midpoint can be modulated by point mutants anywhere along the signal transduction pathway between the sensor and the autokinase. The only requirement is that the substitution has an effect on the internal equilibrium constant for the kinase-promoting versus the phosphatase-promoting conformations of the domain that houses the mutant. So long as the domains are tightly coupled, then an n-fold change in the internal equilibrium will translate to an n-fold change in the midpoint of the overall dose-response curve. Moreover, as the couplings ⍺_1_ and ⍺_2_ become less strong, the magnitude of the shift in the dose response curve is decreased. Thus, it is not necessary to change the binding interactions with the metal ions to affect changes in the ligand sensitivity of the system which provides the system a wealth of opportunities to tune sensitivity.

The fractional change in the kinase activity that can be achieved upon saturation of the ligand-binding sites is a second factor, which ranges with the requirements of a system. For example, the change in transcriptional response in PhoQ is modest, reaching about a factor of 5 to 20-fold change, while other two-component systems such as VirA have a dynamic range as large as 10^5^ (52,55–57). For simple systems that respond to a single ligand, the maximal response is defined by the ratio of the intrinsic affinity of the sensor for the ‘kinase-on’ versus ‘kinase-off’ conformations (K_dON_/K_dOFF_). Mutations that decrease the coupling attenuate the maximal response, and the system becomes decoupled when ⍺_1_ or ⍺_2_ reaches 1. The maximal response in absolute terms is another factor, which depends on the kinetic efficiency of the underlying autokinase domains. When untethered from the membrane, some autokinase domains show large increases activity, so the role of the remainder of the protein can be seen as a negative regulation. Indeed, we find that K_AK_ is significantly less than 1, and this value can be positively modulated by some mutants that reach transcriptional levels somewhat greater than WT for PhoQ. In summary, there is a diversity of mechanisms that nature can call upon to alter the activity of HKs, as illustrated in a relatively small sampling of the 35 mutants studied here.

## Discussion

It has been appreciated for several decades that the effector domains of multi-domain signaling proteins can produce responses that are either more potentiated or diminished relative to the change in state of sensory domains that drive these responses. S J Edelstein and J P Changeux in seminal work coined the terms “hyper-responsive” and “hypo-responsive” to define this uncoupled behavior between domains, which has since been examined in other classes of multi-domain signaling proteins such as GPCRs (58). In this work, we examine the coupling behavior between the sensor and effector domains of transmembrane bacterial sensor histidine kinases and possible roles of modularly inserted signal transduction domains in optimizing this coupling behavior using a model gram negative HK, PhoQ. We find that the intervening HAMP signal transduction domain is necessary to assemble an overall bistable histidine kinase from Mg^2+^-sensor and autokinase-effector domains that are too biased to one signaling state (‘kinase on’ state). This is accomplished by strongly coupling the thermodynamically preferred state of the HAMP to the disfavored ‘kinase off’ signaling states of sensor and autokinase, ameliorating these otherwise strong equilibria such that the overall assembly is bistable and significantly modulated by ligand binding. Thus, the HAMP does more than transmit the response; it instead serves to tune the ligand-sensitivity amplitude of the response.

Evolutionarily, the insertion of signal transduction domains in HKs could allow for the facile modulation of the intrinsic equilibria of sensor and effector domains and their coupling behavior, which may be more difficult to alter through the direct mutation of these domains themselves. The sequence and subsequent structures of sensors and autokinase domains are subject to many evolutionary constraints, be it the specificity and affinity for ligands in sensor domains, the specificity for membrane homodimerization of HKs (59), or the cognate specificity for response regulator (13,60–63) and the ability to inhabit and switch between the various conformations required for a full catalytic cycle in the autokinase domain (15). Furthermore, most two-component systems feature multiple accessory protein components involved in sensing, feedback regulation and cross-talk with other signaling systems, which add evolutionary constraints to these domains (64). In the closely related class of chemotaxis proteins, the analogous transmembrane protein is also subject to extensive covalent modifications that modulate activity. When all these evolutionary activity and specificity considerations are met, the resulting domain may not be ideally bi-stable in isolation. Indeed, in PhoQ, we find that both sensor and autokinase highly prefer the ‘kinase-on’ state, and therefore cannot be allosterically connected to make an overall bistable protein capable of being converted to the ‘kinase-off’ state by Mg^2+^ binding. The presence of one or more signal transduction domains allows for 2 advantageous considerations for producing and finetuning overall HK bistability; the thermodynamic stability of the signal transduction domain can be used to preferentially stabilize or destabilize a given signaling state of sensors or autokinases indirectly through allosteric coupling, and the strength and even direction of coupling can be easily modulated through mutations at the domain junction, rather than mutations that may alter the core functions of the sensor/autokinase themselves.

The latter phenomenon is especially potent in the context of alpha-helical coiled-coil connections between domains of HKs, in which a drastic change in coupling or thermodynamic stability can be caused by minor sequence insertions, deletions and alterations due to the highly regular and cooperative nature of coiled-coil stabilizing interfaces. We have shown that the insertion of a stretch of glycine residues is sufficient to completely uncouple domains. On the other extreme, a well folded coiled coil junction can create strong allosteric coupling due to the cooperative folding and stability of such a motif. A range of stabilities can be achieved by various means, including the insertion or deletion of one or more residues to disrupt the canonical heptad pattern of hydrophobic residues of the dimeric core of the protein, as is often observed in the conserved S-helix motif, which connects HAMP to autokinase domains in HKs (65). Schmidt *et. al.* (66) showed that crystal structures of cytoplasmic domains in different conformations accommodate the structural deviations of these S-Helix sequence insertions by diffusing the strain over different lengths of the proximal alpha-helical core. These different “accommodation lengths” could be analogous to the different strengths of allosteric coupling depending on the signaling states of the adjacent domains in our equilibrium signaling model. We also find conservation of glycine motifs and helix-disrupting proline residues in the juxta-membrane regions of chemotaxis proteins and HKs respectively (51,67,68), which hint at the significant modulation of allosteric coupling strength by the alteration of helical and coiled-coil geometries. In some systems, domains are even segregated to entirely different proteins, in which case the strength of the protein-protein interaction between components can be altered to vary allosteric coupling. These are all evolutionarily accessible solutions to fine-tune the function of a histidine kinase.

Finally, this evolutionary argument may also explain the lack of a parsimonious structural mechanism for signal transduction, even in HKs with a specific domain architecture. Although this problem is largely exacerbated by the dearth of multi-domain structures of HKs in various signaling conformation, several signaling hypotheses have been put forward regarding the structural mechanism for signal transduction in HKs, particularly in HAMP domains. These include the gear-box mechanism (AF1503, Aer2 multi-HAMP)(69), Piston mechanism (Tar) (70, 71), Scissoring mechanism (Tar, BT4663, PhoQ) (72–74), Orthogonal displacement mechanism (HAMP tandems, Tar) (75–77) and the dynamic HAMP mechanism (Adenylate cyclase HAMP) (78–80). A recently elucidated set of structures of the sensor, TM and signal transduction domains of NarQ remains the only representative of a multidomain transmembrane structure of an HK containing a signal transduction domain, and again shows a rigid-body bending transition of the HAMP domain about the conserved N-term Proline between apo- and holo-states of the sensor(81).

It may be that signal transduction mechanisms in HKs are as varied as their modular architecture, and many structural transitions could account for the underlying concern in signaling, which is the allosteric modulation of multi-state equilibria of adjacent domains in response to structural transitions caused by a sensory event. Indeed, the only requirement for signal transduction is a series of domains with two states that either favor or disfavor the kinase state, and a means to transmit the information between the states. Helical connections between domains provide efficiently coupling, but the conformational changes within the domain need not be obligatorily the same for different domains. Additionally coupling can involve tertiary contacts, which can be used in conjunction with or instead of helical connections. Interestingly, the observation that PhoQ has a weakly HAMP-coupled ‘kinase-on’ state and a strongly HAMP-coupled ‘kinase-off’ state has been posited before, albeit in the context of a hypothesized tertiary contact between the membrane-distal portion of HAMP helix-1 and a loop in the autokinase domain (54). The idea that autokinase domains intrinsically have high-kinase activity and are subsequently inhibited by strong coupling to up-stream domains and the further stabilization of these inhibitory conformations by ligand-binding warrants examination as a generalizable signaling mechanism for histidine kinases.

## Materials and Methods

### Materials

BW25113 and HK knockout strains were obtained from the Keio collection.

TIM206 (*E. coli* Δ*phoQ*, p*mgrB*::LacZ) was obtained from Tim Mayashiro (Goulian lab)

pTrc99a (GenBank # M22744)

pSEVA311 (GenBank# JX560331) was developed by the de Lorenzo lab and was a gift from the European Standard Vector Architecture consortium.

Brilacidin was a gift from Polymedix Inc. N-ethylmaleimide (NEM, Sigma)

Tris-Acetate gels (Thermofisher Scientific)

Anti-PentaHis antibody (Thermofisher Scientific)

### Methods

#### Cloning

PhoQ mutants were cloned into the pTrc99a plasmid MCS by restriction cloning. Point mutations were made by quick-change mutagenesis and confirmed by sanger sequencing. Hybrid HK-gene reporter plasmids were built in pTrc99a plasmid by introducing a c-terminally 6x His-tagged HK construct into the IPTG inducible MCS, and the mCherry reporter sequence downstream by Gibson cloning. Sequences of reporters are available in supplementary methods. Gly_7_ disconnections and point mutations were introduced by a blunt-end ligation strategy and confirmed by sanger sequencing.

#### Growth of PhoQ constructs

For each biological replicate, an isolated colony of TIM206 (genotype: Δ*phoQ*, p*mgrB*::LacZ) containing various pTrc99a-*phoQ* constructs was grown overnight at 37 ℃ in MOPS minimal media + 50 µg/mL AMP and 1 mM MgSO_4_. These overnight cultures were then diluted 50x into 1 mL MOPS media + 50 µg/mL AMP and 1 mM MgSO_4_ and grown at 37°C for 2 hours. These cultures were further diluted 500X into 30 mL MOPS minimal media + 50 µg/mL AMP containing 0.1, 0.4, 1.6, 6.4 and 25.6 mM MgSO_4_, and grown for at least 5 hours such that the density of the culture reaches log-phase (OD_600_ = 0.2 – 0.8). 500 µL of culture is removed for evaluating beta galactosidase activity, while the remaining culture is used for western analysis. Two constructs (A225F, Y265A Gly_7_ 260/261) required induction with 10 µM IPTG during growth for observable levels of membrane inserted PhoQ by western blot.

#### Beta galactosidase activity

500 µL of PhoQ culture was combined with 500 µL of 1x Z-buffer + 40 mM beta-mercaptoethanol, 25 µL of 0.1% SDS in water, and 50 µL of chloroform in a glass culture tube and vortexed for complete lysis. The lysate was then prewarmed to 37°C in a standing incubator before addition of ONPG substrate. 0.25 mL of prewarmed 4 mg/mL ONPG in 1x Z-buffer + bMe was added to the lysate to initiate hydrolysis, which was then quenched with the addition of 500 µL of 1M Na_2_CO_3_ after variable incubation periods. The quenched hydrolysis was then centrifuged to remove any cell debris, and absorbance at 420 nm and 550 nm was measured in triplicate using a Biotek synergy2 plate-reader with pathlength correction. Miller units were calculated as follows:

Miller units = 1000*(OD_420_ – 1.75*OD_550_)/(OD_600_*dilution factor*incubation time in min)

#### Membrane fraction isolation

30 mL of PhoQ culture was centrifuged at 4°C for 20 minutes to collect a cell-pellet. This cell-pellet was immediately frozen in liquid nitrogen and stored at −80°C until analysis. Frozen pellets were first thawed, suspended and incubated on ice with 500 µg/mL N-Ethylmaleimide (NEM) and 1 mg/mL lysozyme in 50 mM TRIS buffer, pH 8, for 1 hour. Cells were then lysed by 30 seconds of tip-sonication. Lysed cells were then centrifuged at 16000xg for 10 minutes to remove cell debris. Membrane was isolated from the supernatant by further centrifugation at 90,000xg for 10 minutes. Membrane pellets were then resuspended in 1X LDS loading buffer containing 8M Urea and 500 mM NEM, boiled at 95°C for 10 minutes and analyzed by western blot.

#### Monomer and dimer quantification by western blot

Samples were first separated on 7% TRIS-SDS gels by electrophoresis at 200V for 70 min, and then transferred onto nitrocellulose membranes by dry transfer (iBlot2). Membranes were then blocked using 1% BSA in TBS-t buffer (20 mM Tris, 2.5 mM EDTA, 150 mM NaCl, 0.1% Tween-20), probed using an anti-pentaHis HRP antibody, and visualized using luminescent ECL substrate on a BioRad imager. Bands corresponding to PhoQ monomer and dimer were quantified using Image-J software to yield a crosslinking efficiency between 0 and 1.

#### Measuring activity of CpxA, BaeS

HK constructs were cloned into the MCS of pTrc99a plasmid, and the associated fluorescent reporter gene was cloned downstream. For the CpxA reporter plasmid, the response regulator CpxR, was also cloned into the MCS and transformed into AFS51 strain (Δ*cpxA*Δ*pta*::Kan p*cpxP*::GFP) by heat shock transformation. For BaeS, the response regulator BaeR, was cloned into an additional plasmid, pSEVA331 under an IPTG inducible promoter and both plasmids were transformed into a Δ*baeS*Δ*cpxA* double KO strain by heat shock transformation. Cultures were started by diluting overnights 200-500 folds into fresh LB + 50 µg/mL AMP media and allowed to grow to mid-log phase (OD_600_ = 0.4 – 0.6) before analysis by flow cytometry. The responsiveness of *cpxP* reporter was confirmed by treating log-phase cultures with 2 µg/mL Brilacidin for 1.5 hours before analysis. Expression of HKs was confirmed by western analysis using the c-terminal 6x His-tag for quantification.

#### Flow cytometry

LB cultures at mid-log phase were diluted 20x into 1x PBS buffer and 20,000 cells gated by forward and side-scatter were evaluated for GFP fluorescence (p*cpxP*::GFP; Ex. 488 nm, Em. 515 nm) or mCherry fluorescence (p*spy*::mCherry, Ex. 488 nm, Em. 620 nm) per sample on a BD FACS caliber instrument. Sample average fluorescence and standard error were determined by standard analysis using Flo-Jo software.

#### Data Fitting

For data fitting, only data-sets in which kinase activity and sensor crosslinking have been determined simultaneously from the same samples at all 5 concentrations of Mg^2+^ were included in analysis. This resulted in Kinase active and sensor crosslinking competent states are partitioned to generate expressions dependent on [Mg^2+^] as the lone variable as shown below. The parameters are then fit globally across all datasets, except for those accounting for the perturbation of a mutant/ Gly_7_ disconnection, which are fit locally. Locally fit parameters are kept identical between replicates or additive mutations.

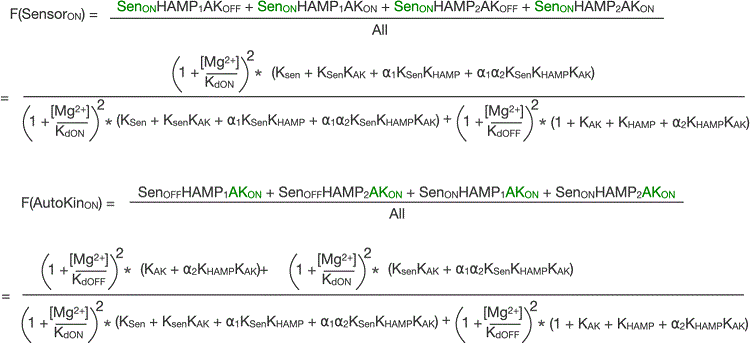

To ensure equal weights in global fitting, the activity data was scaled by a factor of q = (mean of activity data) / (mean of %crosslink data). The crosslinking data and refactored activity data (Activity / q) were then globally fit to a 3-state allosteric model. Each of 56 datasets (including replicates) was fit by a combination of global and local parameters, described in **Table 2**. Global parameters were shared between replicate datasets as well as datasets of mutations that were functionally similar. A total of 62 parameters (global and local, **Table 2**) were optimized using the python code found in the supplement (phoq_fit_local_global.py), from many rounds of fitting starting with random initial conditions (125,000 independent fits). Error analysis of the best-fit parameters (minimized sum of squares of residuals) was performed through bootstrapping of residuals with replacement to calculate confidence intervals, as well as residual sweep analyses (see below). To create synthetic bootstrapped datasets, we chose residuals at random with replacement and added these residuals to the activity and %crosslink values from the optimum fit. For each synthetic dataset, parameters were re-optimized, starting from initial values taken from the optimum fit. Out of 10,000 generated datasets 3061 fits were determined to have converged. The optimization process was considered converged when the cost function F did not change considerably (dF < ftol * F, with ftol = 1e-8, i.e., convergence criterion 2 from Scipy least_squares). Histograms of these bootstrapped parameter values show the spread in possible values due to errors in the fit (**Figure 7c** and **Figure 8 figure supplement 1**). Analysis of the bootstrapped parameter distributions showed correlations between the globally fit parameters S and K_AK_ (**Fig. 7 figure supplement 1**).

**Table 2.**
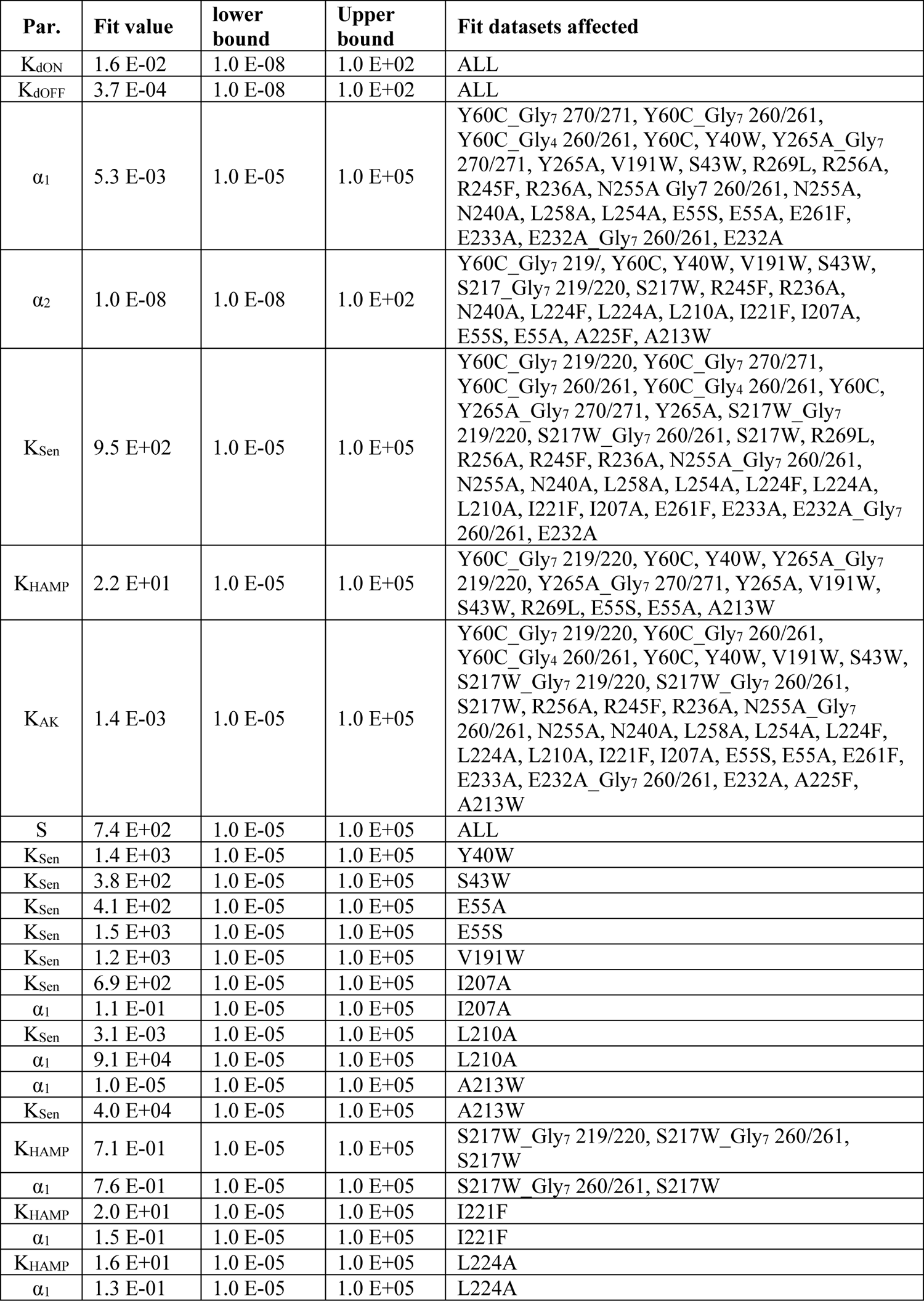
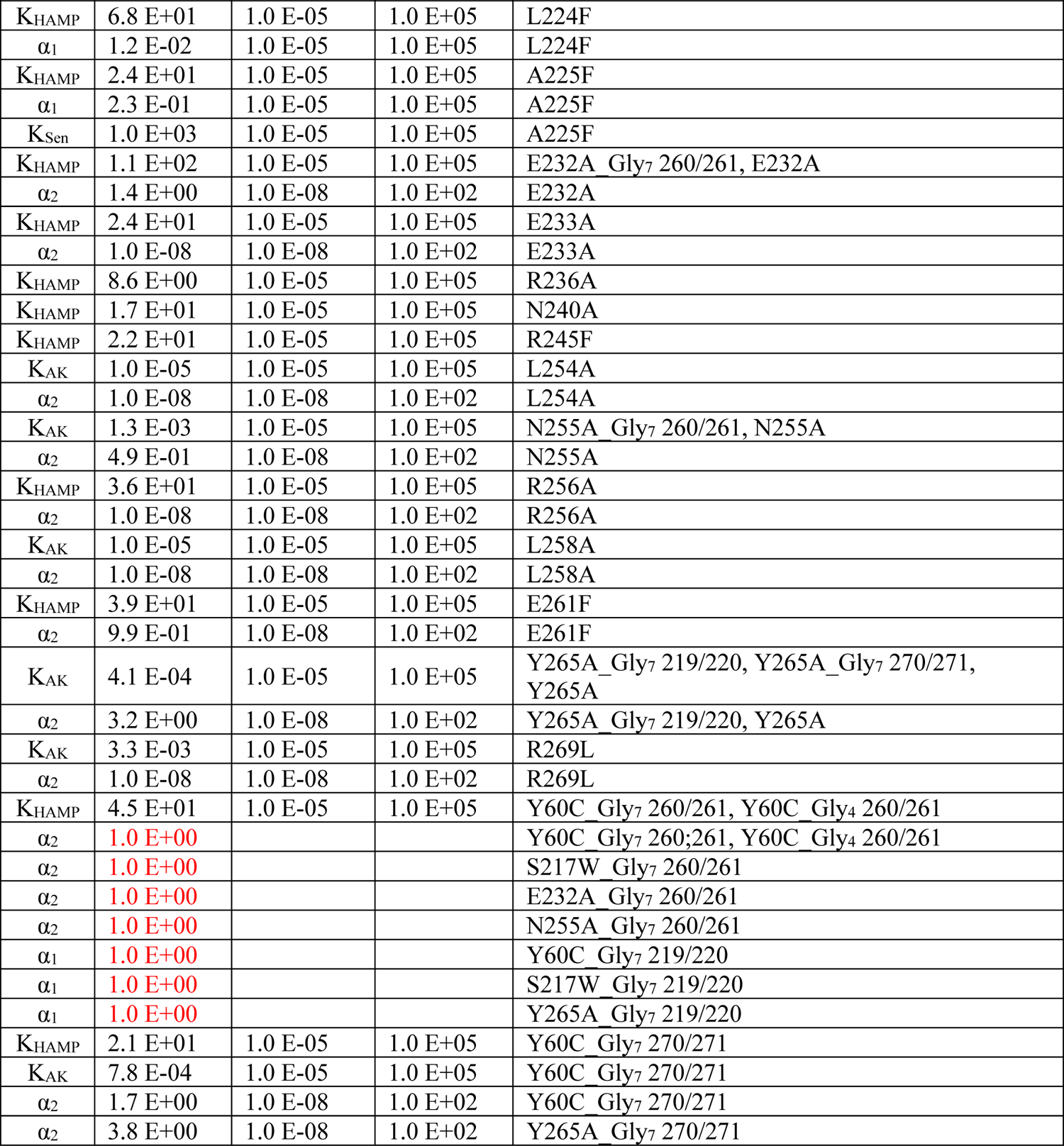
parameters used in fitting

We also performed a residual sweep analysis to assess the quality of the fit in response to changes in a single parameter value, with all other parameters held fixed. For residual sweep analysis, all but one of the parameters were fixed to their optimum values, and the variable under analysis was swept across its allowed numerical range, after which the sum of squares of residuals was calculated. The sum of squares was then plotted as a function of the parameter’s numerical value (**Figure 7C** and **Figure 8 figure supplement 1**). Code to reproduce the fits and plots is given in the comment section at the bottom of the supplement python scripts (phoq_fit_local_global.py, phoq_fit_local_global_ipython.py, and phoq_fit_ci_local_global.py). Scripts to run the fitting on the UCSF Wynton High Performance Computing cluster can also be found in the supplement (phoq_fit.job and phoq_fit_ci.job).

#### Choice of locally varied parameters

mutations contained entirely within a given domain are allowed to vary the intrinsic equilibrium of that domain only. Mutations within 1 heptad of a domain-domain junction (219/220 for sensor/HAMP, 260/261 for HAMP/autokinase) are also allowed to vary the equilibrium constant of the domain they reside in, as well as the coupling constant between the two domains. Exceptions to this rule include A225F, which was additionally allowed to vary the K_Sen_ parameter, along with K_HAMP_ and ⍺_1_ parameters which would normally be varied. Given the poor fit to this mutant, we hypothesized that the disruption of inserting a large Phe sidechain in place of an alanine may propagate into the preceding transmembrane region. Similarly, we allowed K_AK_ to float locally for L254A, N255A and L258A, which resulted in better fits as discussed in main text. Finally, ⍺_2_ was allowed to float for E231A, and E232A, which have been hypothesized in previous work to directly couple to the autokinase domain via a salt-bridge to an arginine residue in the autokinase (54).

## Acknowledgements.

Funding: We acknowledge research support from a grant from NIH (R35 GM122603).

